# CglB adhesins secreted at bacterial focal adhesions mediate gliding motility

**DOI:** 10.1101/2020.07.22.216333

**Authors:** Salim T. Islam, Laetitia My, Nicolas Y. Jolivet, Akeisha M. Belgrave, Betty Fleuchot, Gael Brasseur, Laura M. Faure, Gaurav Sharma, David J. Lemon, Fares Saïdi, Jean-Bernard Fiche, Benjamin P. Bratton, Mitchell Singer, Anthony G. Garza, Marcelo Nollmann, Joshua W. Shaevitz, Tâm Mignot

**Author notes:** co-corresponding authors Tâm Mignot, Salim T. Islam Phone: (+1).

## Abstract

The predatory deltaproteobacterium *Myxococcus xanthus* uses a helically-trafficked motor at bacterial focal adhesion (bFA) sites to power gliding motility. Using TIRF and force microscopy, we herein identify the integrin αI-domain-like outer-membrane (OM) lipoprotein CglB as an essential substratum-coupling protein of the gliding motility complex. Similar to most known OM lipoproteins, CglB is anchored on the periplasmic side of the OM and thus a mechanism must exist to secrete it to the cell surface in order for it to interact with the underlying substratum. We reveal this process to be mediated by a predicted OM β-barrel structure of the gliding complex. This OM platform was found to regulate the conformational activation and secretion of CglB across the OM. These data suggest that the gliding complex promotes surface exposure of CglB at bFAs, thus explaining the manner by which forces exerted by inner-membrane motors are transduced across the cell envelope to the substratum; they also uncover a novel protein secretion mechanism, highlighting the ubiquitous connection between secretion and bacterial motility.

## Introduction

Directed surface motility of cells from all biological kingdoms involves highly-dynamic cell–substratum interactions. In eukaryotic cells, this process involves the engagement and activation of surface-exposed integrin(-like) adhesins, directionally transported by molecular motors (myosin) via integrin coupling to the internal cytoskeleton (actin)^1^. For metazoan organisms, nascent integrin adhesions to the extracellular matrix (ECM) lead to integrin nucleation and the formation of large eukaryotic focal adhesion (eFA) sites; these assemblies remain fixed-in-space relative to a translocating cell, promoting local traction, transduction of motor forces, and cell translocation^2^. Such surface motility is not restricted to eukaryotic cells. Long known to move on surfaces in the absence of outward appendages (such as flagella or type IV pili) via a process termed “gliding motility”^3^, individual cells of the Gram-negative predatory deltaproteobacterium *Myxococcus xanthus* were shown to utilize a trans-envelope multi-protein Agl–<u>Gl</u>iding transducer (Glt) complex (**Fig. 1A**) to power gliding motility^4, 5^. In motile cells, these complexes associate at the leading pole and move directionally in the bacterial inner membrane (IM) toward the lagging cell pole, following a right-handed helical trajectory^6–8^ (**Fig. 1A**). These rotational movements likely probe the substratum beneath gliding cells, leading to immobilization of the Agl–Glt complex at fixed bacterial focal adhesion (bFA) sites as well as cell translocation via left-handed rotation of the bacterium around its long axis^8^ (**Fig. 1A**).

**Figure 1:**
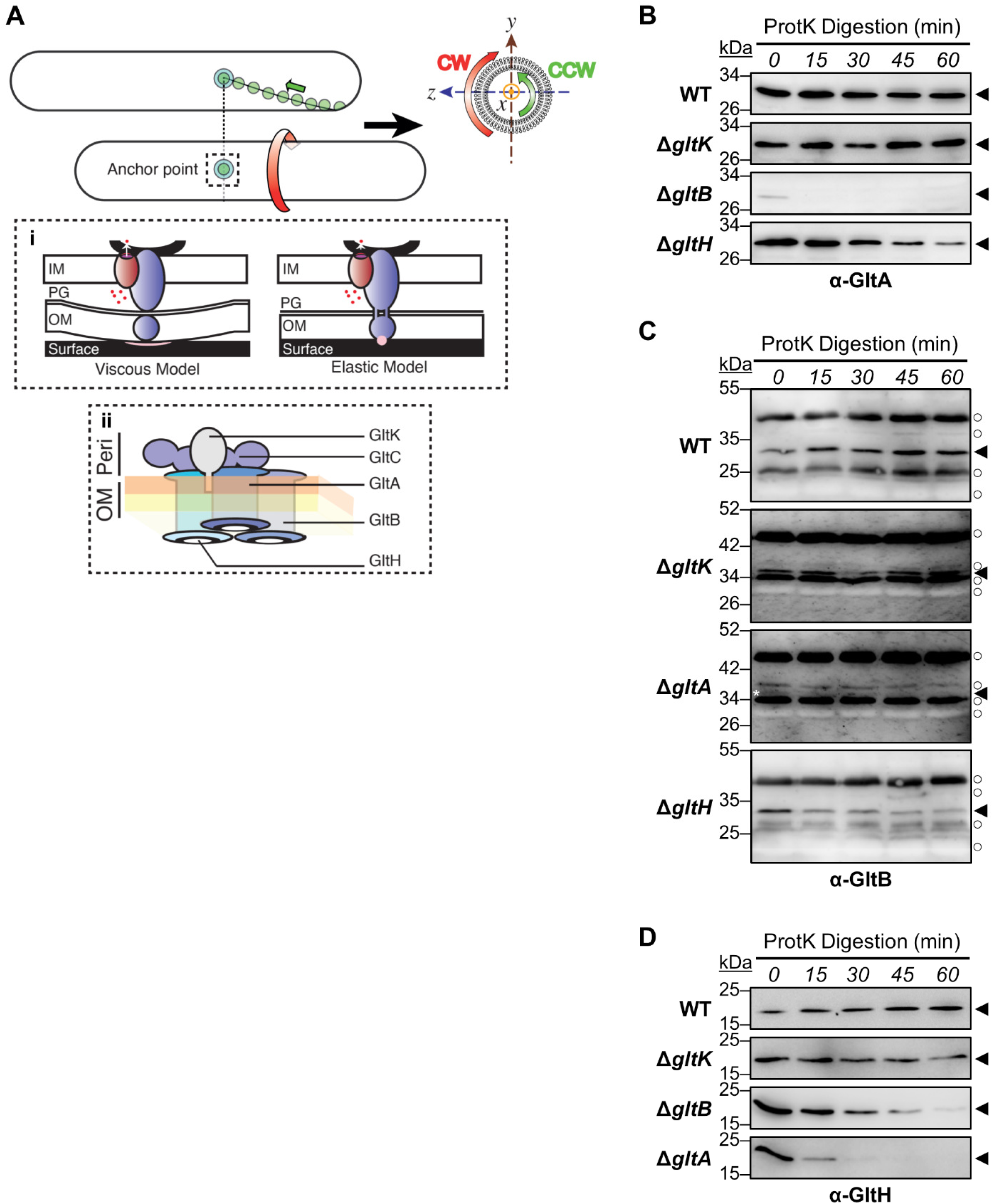
Evidence of an Integral OM Complex Formed by GltA, B, and H. **A)** Gliding motility mediated by bFAs in *M. xanthus*. Following their assembly at the leading pole, motility complexes move toward the lagging cell pole in a counter clockwise (CCW) rotational trajectory. Clockwise (CW) and CCW directionalities are defined by observing the cell cylinder from the leading pole in the *y,z* plane. When the complexes interact with the substratum, they form bacterial focal adhesion (bFA, concentric circles) sites and propel rotational movements of the cell. *Panel (i)*: bFAs are formed according to two possible mechanisms: In the viscous interaction model, the periplasmic complex accumulates at bFAs and pushes against the elastic peptidoglycan (PG) to create cell envelope deformations at bFAs and thus create viscous interactions with the substratum. The function of the outer-membrane (OM) complex is not accounted for in this model. In the elastic model, the periplasmic complex establishes transient interactions through the PG, contacting the OM complex which itself interacts with the substratum via an unknown adhesive molecule (*pink circle*). *Panel (ii):* Proposed Glt OM platform based on previous reports and this study. The OM localization of GltA, GltB, GltH, GltC and GltK is based on structural bioinformatic as well as fractionation analyses presented here and elsewhere^4, 13, 14^. The integral association of GltA, B and H is based on bioinformatic and Proteinase K accessibility assays in this study and another report^13^. Direct GltA–GltB, GltA–GltC, and GltB–GltC interactions were already biochemically demonstrated by pull-down assays^13^. The connection with GltH is further indicated from results reported in this study. The periplasmic leaflet (*orange*) and outer leaflet (*yellow*) of the OM are indicated. Legend: Peri, periplasm. Western immunoblot of **B)** GltA, **C)** GltB, and **D)** GltH susceptibility to digestion by Proteinase K in OM-module Agl–Glt mutant strains. Digestion aliquots were removed at 15-min intervals and TCA-precipitated to stop digestion. Legend for Panels B, C, and D: ◄, full-length protein; ○, loading control (non-specific protein band labelled by the respective α-GltA/α-GltB/α-GltH pAb).

At the molecular scale, the mechanisms that promote force transmission and adhesion at bFAs are not known. The IM components of the motility complex assemble at the pole on a cytoplasmic scaffold formed by bacterial actin MreB, the Ras-like protein MglA and the coiled-coil AglZ protein^9, 10^. The IM complex contains the molecular motor AglRQS, a TolQR/ExbBD/MotAB-like channel that uses the proton gradient formed across the bacterial IM to energize long-range movements of the IM complex in the bacterial envelope^11^. However, the manner in which these intracellular motions are coupled to the substratum in order to propel the cell is unknown. One hypothesis states that trafficking motor units deform the peptidoglycan meshwork in the periplasm, propagating surface wave deformations and viscous interactions between the outer membrane (OM) and the substratum (**Fig. 1A, panel i**). However, observations and mechanical modelling of cell–cell collision events suggest that interactions between a gliding cell and the substratum are elastic in nature, consistent with localized adhesion points and the existence of an anchored adhesin^12^ (**Fig. 1A, panel i**). In the cell envelope, the IM motor moves by establishing transient contacts with a complex of Glt proteins localized in the OM (herein called the OM platform, see below), linked via periplasmic domains of retractile proteins that traverse the peptidoglycan meshwork^8^ (**Fig. 1A**). When in contact with the substratum, these motions tether the OM platform at bFAs, which is proposed to create local adhesions and movement of the cell (**Fig. 1A**). Consistent with localized adhesion at bFA sites, gliding cells abandoned patches of OM motility-complex proteins on the substratum at positions formerly occupied by bFA sites^8^. However, given that the OM platform is required for motor movements, it could not be assessed whether the OM platform is directly responsible for adhesion to the substratum^3, 8^ and thus, the manner of adhesion, a core aspect of the motility mechanism, remained unaddressed.

The proteins GltA, GltB, GltH, and GltK constitute the proposed OM platform (**Fig. 1A, panel ii**). While the exact structure of this platform is not known, GltA, GltB, and GltH are all localized in the OM and are predicted to form integral OM β-barrels^3, 4, 8, 13, 14^ (**Fig. 1**, **Supplementary Fig. S1A-C**). GltA and GltB interact with each other, as well as with GltC present in the periplasmic leaflet of the OM^13^ (the function of GltC is not addressed herein). GltK and GltH interactions have yet to be defined but data presented in our investigation suggest that these proteins are functionally associated with the OM platform. Herein, we also identify the OM lipoprotein CglB^15^ as a main substratum-coupling adhesin of the Agl–Glt gliding motility apparatus associated with the OM platform. Specifically, the results support the presence of a protein assembly formed by GltA, B, H, and K that regulates CglB surface exposure, secreting it to the cell surface upon stimulation of the motility complex at bFAs.

## Results

### GltA, B, and H form a putative integral OM protein complex

The Glt OM platform contains predicted OM-spanning β-barrel proteins (**Fig. 1**, **Supplementary Fig. S1A-C**) and thus is unlikely to mediate direct substratum-anchoring of the motility complex. To probe the integral OM nature of these proteins, we tested the sensitivity of GltA, GltB, and GltH in the OM to digestion by Proteinase K. In this assay, proteins that are largely surface-exposed are rapidly digested by addition of exogenous protease, unlike proteins that are protected by the membrane environment^13, 16^. As expected for predicted integral OM proteins, GltA, GltB, and GltH were not readily susceptible to digestion by Proteinase K at the surface in WT cells (**Fig. 1B,C,D**). To probe the presence of an OM-platform protein complex, the Proteinase K susceptibility of a given constituent was tested in the absence of a different OM-platform protein; the rationale was that cell-surface topology for various OM-platform proteins could be altered due to a disrupted interaction network resulting from the missing platform component. As previously detected^13^, neither GltA nor GltB was stable in a mutant background lacking the other, as reflected by the lower levels of GltA in Δ*gltB* cells and GltB in Δ*gltA* cells (compared to WT) (**Fig. 1B,C, Supplementary Fig. S1D**). This is consistent with the demonstrated interactions between these integral OM proteins^13^. Absence of GltK did not alter Proteinase K susceptibility of GltA, GltB, or GltH, indicating that this periplasm-facing OM lipoprotein^13^ does not affect the topology of the OM-platform proteins (**Fig. 1B,C,D**). However, the absence of GltH rendered GltA and GltB more Proteinase K-sensitive (**Fig. 1B,C**). Similarly, GltH was more sensitive to Proteinase K digestion in the absence of GltA or GltB (**Fig. 1D**). The potential for these Proteinase K-susceptibility changes to be due to general OM defects in the respective mutants is unlikely as none of the mutant strains demonstrate obvious membrane and/or cell fitness defects^13^. These (and previous^13^) data thus support the existence of an integral OM complex formed by GltABH proteins (**Fig. 1A, panel ii**). Functional analysis of the role of the OM platform in regulating gliding-complex adhesion to the substratum was subsequently evaluated below.

### CglB, a predicted α-integrin-like protein, is a candidate motility adhesin

We next searched for a candidate adhesin that might interact with the OM platform. The CglB protein is an ideal candidate because it is essential for efficient gliding motility^4, 15, 17, 18^ (**Supplementary Fig. S2A**), localizes to the bacterial OM^14^, and is pulled down as prey with various gliding complex members via GST chromatography using Glt-protein bait^5^. The *cglB* gene also co-occurs with those encoding the Agl–Glt machinery in bacterial genomes, supporting a functional link (**Supplementary Fig. S3**). Along with all Agl–Glt gliding apparatus components, we also found CglB to be required for “polymertropism” (**Supplementary Fig. S2D**), as determined by the ability of *M. xanthus* to preferentially spread in the direction of aligned substratum polymers^19–21^. This suggests that these proteins may play a role in probing the physical properties of the substratum at bFAs, similar to eukaryotic integrins that interact with the substratum and sense its physical properties at eFAs^22–24^.

Fold-recognition analysis of CglB indicated structural analogies with numerous metazoan α-integrins and structurally-similar Apicomplexan parasite gliding motility adhesins MIC2 and TRAP (from *Toxoplasma* and *Plasmodium*, respectively)^25, 26^ (**Supplementary Table S2**). The top bacterial match was the αI/αA-like GBS104 adhesive tip pilin^27^ from *Streptococcus agalactiae* (**Supplementary Table S2**). Integrins typically possess an α subunit in which a C-terminal domain of seven repeating beta blades constitutes a beta-propeller domain^28^. Of the 18 human α-subunit variants, half possess an additional module (termed αI or αA) in-between the 2^nd^ and 3^rd^ propeller blades containing a VWA domain^29^, characterized by a Rossmann fold with multiple α-helices shielding an interior β-sheet^30^. VWA domains typically function in cell adhesion and can be found in numerous proteins of the eukaryotic ECM. Generation of a CglB tertiary structure model, combined with analysis of evolutionarily-coupled amino acid positions within the protein, confirmed that CglB likely adopts the above-described αI/αA structure containing a VWA domain (**Fig. 2A, Supplementary Fig. S1E**). In particular, the VWA domain contains a predicted MIDAS (<u>m</u>etal ion-<u>d</u>ependent <u>a</u>dhesion <u>s</u>ite) motif, a discontinuous structural feature (Asp-x-Ser-x-Ser…Thr…Asp). At this site, the coordination of a divalent metal ion (e.g. Mg^2+^/Mn^2+^/Ca^2+^) induces structural changes in VWA domains upon ligand binding that stabilize this adhesive domain in a high-affinity state for the ligand^31^. Incidentally, CglB-dependent gliding by individual cells on a hard substratum (**Fig. 2C**) was also found to require the presence of divalent cations (**Supplementary Figure S2G**). Highly-conserved potential MIDAS residues in CglB map to D56, S58, S60, T182 and D211 (**Fig. 2A, inset**). We tested the functional significance of the putative CglB MIDAS motif in a complementation assay, ectopically expressing CglB_WT_, CglB_D56A_, and CglB_S58A_ from the native *cglB* promoter. Contrary to Δ*cglB* cells in which CglB_WT_ expression restored motility, CglB_D56A_ was stably expressed (unlike CglB_S58A_) but unable to complement gliding deficiency (**Supplementary Fig. S2E, F**).

**Figure 2:**
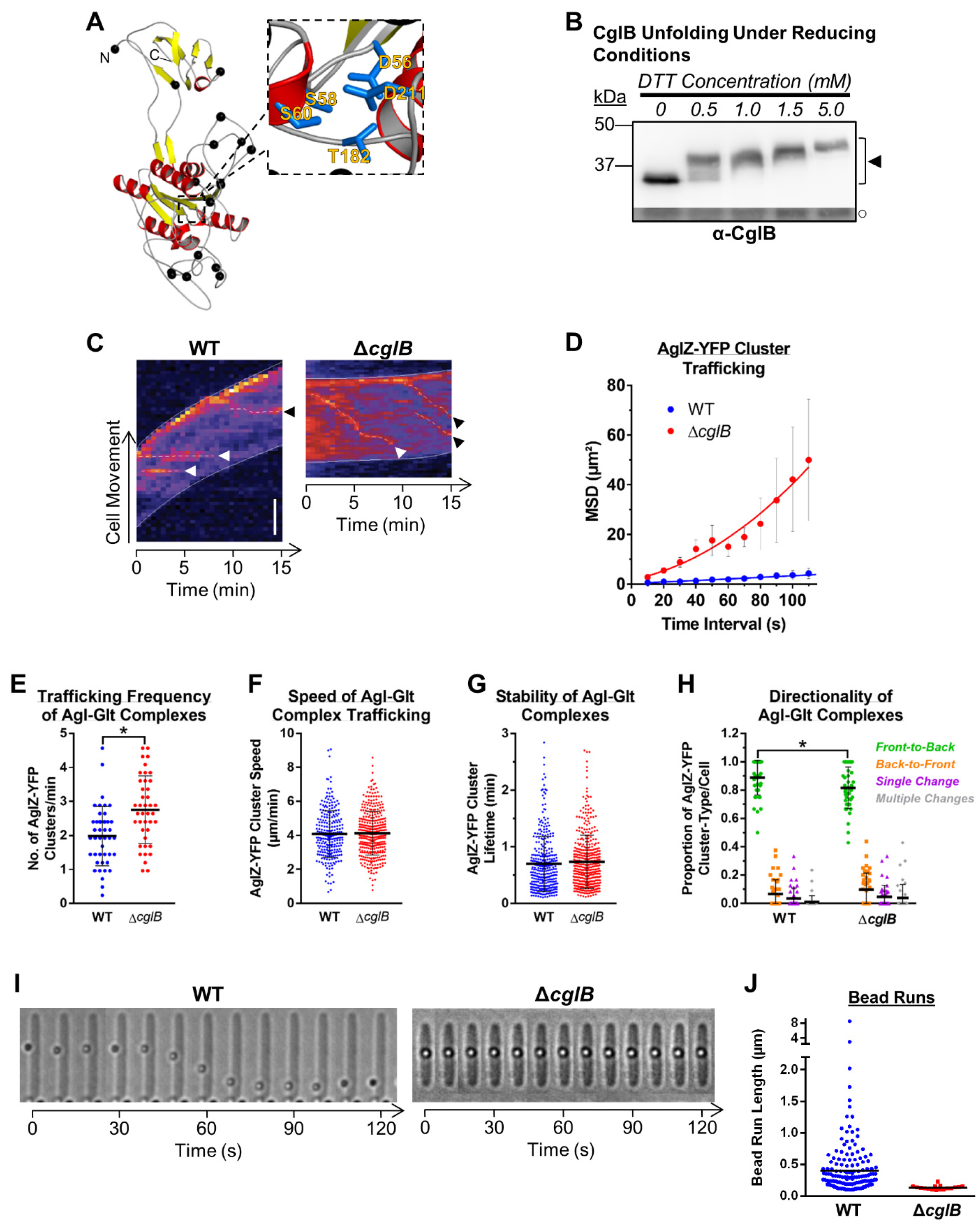
CglB, a protein with a potential α-integrin-like fold, is essential for motility complex adhesion to the substratum. **A)** Tertiary structure homology model of CglB. The N- and C-terminus of the protein are indicated. Colour code: *yellow*, β-strands; *red*, α-helices, *black spheres*, Cys residues. Dotted line denotes a magnified view of the MIDAS motif containing residues D56, S58, S60, T182, D211. See also **Supplementary Figs. S1E, S2E, F** and **Supplementary Table S2**. **B)** α-CglB Western blot of WT whole-cell extracts treated with increasing concentrations of DTT to break disulphide bonds. Legend: ◄, full-length CglB; ○, loading control (non-specific protein band labelled by α-CglB pAb). See also **Supplementary Fig. S2B**. **C)** Kymograph in WT vs. Δ*cglB* cells indicating AglZ-YFP cluster position over time (hashed lines). Scale bar: 2 µm. Legend: white arrowheads, AglZ-YFP clusters followed for their entire lifetime; black arrowheads, AglZ-YFP clusters followed for an incomplete lifetime. See also **Supplementary Fig. S4B**. **D)** Mean-squared displacement (MSD) analysis of AglZ-YFP cluster position tracking in WT (n = 48 clusters) and Δ*cglB* (n = 23 clusters) *M. xanthus* cells. The mean of MSD at each time interval is displayed ± SEM, with a second-order polynomial line fit to each dataset. **E)** Frequency of trafficking Agl–Glt complexes via TIRFM (of AglZ-YFP) on chitosan-coated glass surfaces in PDMS microfluidic chambers for WT (n = 44 cells) and Δ*cglB* (n = 41 cells) strains. The distribution of the two datasets are significantly different (*), as calculated via unpaired two-tailed Mann-Whitney U-test (*p* < 0.05). See also **Supplementary Fig. S4C**. **F)** Speed of Agl–Glt complex trafficking via TIRFM (of AglZ-YFP) on chitosan-coated glass surfaces in PDMS microfluidic chambers for WT (n = 260 clusters) and Δ*cglB* (n = 371 clusters) strains. The distribution of the two datasets are not significantly different, as calculated via unpaired two-tailed Mann-Whitney U-test (*p* > 0.05). See also **Supplementary Fig. S4C**. **G)** Stability of trafficking Agl–Glt complexes via TIRFM (of AglZ-YFP) on chitosan-coated glass surfaces in PDMS microfluidic chambers for WT (n = 333 clusters) and Δ*cglB* (n = 409 clusters) strains. The distribution of the two datasets are not significantly different, as calculated via unpaired two-tailed Mann-Whitney U-test (*p* > 0.05). See also **Supplementary Fig. S4C**. **H)** Directionality of trafficked Agl–Glt complexes via TIRFM (of AglZ-YFP) on chitosan-coated glass surfaces for WT (n = 44 cells) and Δ*cglB* (n = 41 cells) strains. “Front” and “back” are defined as cell poles with high and low AglZ-YFP fluorescence intensity, respectively. Relative to the respective reference WT cluster type, only the distribution of front-to-back clusters were significantly different in the Δ*cglB* cells (*p* < 0.05); all other cluster types did not display distributions different from WT (*p* > 0.05), as calculated via unpaired two-tailed Mann-Whitney U-test. See also **Supplementary Fig. S4C**. **I)** Trafficking of surface-deposited polystyrene beads on *M. xanthus* cells. Images were acquired at 10 s intervals. See also **Supplementary Fig. S4D**. **J)** Length of tracked bead runs > 0.1 µm in *M. xanthus* cells. Images from 10 s intervals were analyzed. See also **Supplementary Fig. S4D**.

An intact VWA domain thus appears essential for CglB stability and single-cell gliding motility. Direct demonstration of metal binding by the MIDAS motif of purified CglB was not possible due to insurmountable purification challenges attributed to the high CglB Cys content (17 out of 416 aa = 4.1%) (**Supplementary Fig. S2B**), with the majority of these residues predicted to localize in flexible loops (**Fig. 2A**). These Cys residues likely form structure-determining intra-protein disulphide bonds, as titration of the reducing agent DTT resulted in a migration shift from faster-to slower-moving CglB-specific bands via SDS-PAGE and α-CglB Western immunoblot (**Fig. 2B**). These data are also consistent with the high sensitivity of single-cell gliding motility to low substratum concentrations of DTT (**Supplementary Fig. S2C**). Taken together, the VWA domain MIDAS motif and intra-protein disulphide bonds are likely important structural determinants of CglB.

### CglB is essential for substratum-coupling of the Agl–Glt machinery

We subsequently investigated the contribution of CglB to surface coupling of the Agl–Glt complex. To probe the role of CglB in bFA formation, we analyzed the dynamics of AglZ-YFP clusters in cells on hard agar for the Δ*cglB* mutant (which stably expresses AglZ-YFP [**Supplementary Fig. S4A**]). AglZ-YFP clusters still appeared in Δ*cglB* cells; however, in marked contrast to WT cells, AglZ-YFP clusters in Δ*cglB* cells were not stationary relative to the substratum but rather moved directionally from one pole to the other (**Fig. 2C,D** and **Supplementary Fig. S4B**). This behaviour was similar to that observed in non-adhered motility complexes^8^. CglB is therefore required to assemble bFAs on hard agar surfaces. The function of CglB is clearly distinct from the OM-platform β-barrel proteins because trafficking AglZ-YFP clusters are not formed in any of the Δ*gltA*/*B*/*H* mutant backgrounds^8^.

In Δ*cglB* cells, trafficking AglZ-YFP clusters move in-and-out of the epifluorescence focal plane as they rotate counter-clockwise around the cell envelope (**Fig. 1A, Supplementary Fig. S4B**), making them difficult to precisely track and thus accurately study their dynamic properties. To resolve these difficulties, we recently developed a TIRFM assay in which *M. xanthus* cells glide in chitosan-coated microfluidic chambers (**Supplementary Fig. S4C**); in this system, trafficking AglZ-YFP clusters are also observable in WT cells due to the suboptimal nature of the chitosan surface for bFA adhesion^8, 32^. Since the depth-of-field in TIRFM is restricted to near the cell–substratum interface, photobleaching is reduced and thus tracking of the trafficking AglZ-YFP clusters near the ventral face of the cell can be performed at high spatio-temporal resolution^8^. On chitosan, Δ*cglB* cells were also non-motile and again, immobilized AglZ-YFP clusters could not be detected (**Supplementary Fig. S4C**). Trafficking AglZ-YFP clusters in Δ*cglB* cells behaved similarly to those in WT cells; although slight effects were observed via TIRFM on the trafficking frequency of AglZ-YFP clusters (from the leading to the lagging cell poles), the trafficking speed and lifetime of AglZ-YFP clusters were unchanged in the absence of CglB (**Fig. 2E-H**). Since AglZ-YFP trafficking reflects the activity of the motility engine, we conclude that CglB does not affect motor activity, but rather its adhesion to the underlying substratum at bFAs.

To test the contribution of adhesive properties by CglB to the tip of the motility complex, we adopted a force microscopy approach; herein, force generation by the motility complex can be directly monitored in live *M. xanthus* cells immobilized atop a semi-solid 0.7% agarose matrix deposited on glass slides^11^. Within this setting, the motility complex cannot propel cells (likely because it cannot adhere) but its activity can transport polystyrene beads deposited by an optical trap directly on the cell surface (**Fig. 2I**). Trafficking gliding-machinery units that collide and recruit such beads move them directionally over long distances^11, 12^ (**Supplementary Fig. S4D**). We therefore tested whether bead transport requires the CglB adhesin. Whereas beads were transported multiple times, at lengths up to ∼8 µm, along the surface of WT cells, such events were absent in Δ*cglB* cells (**Fig. 2J**). This demonstrates that bead recruitment and trafficking requires CglB, consistent with adhesive and force transduction functions for CglB.

Taken together, we conclude that CglB is required for tethering the gliding motility complex to an engaged extracellular motif, be it a solid surface for cell gliding or cargo for transport in immobilized cells. Contrary to the OM-platform proteins GltA, B, and H^8^ (see below), CglB is not required for Agl–Glt-complex assembly and trafficking suggesting that it functions to couple trafficking units to the substratum, as would be expected for an adhesin.

### The Glt OM platform regulates CglB exposure and retention at the cell surface

Our various attempts to localize CglB in gliding cells were unsuccessful due to loss of protein functionality when fused to fluorescent protein or epitope tags (the folding of CglB in the periplasmic space, as well as its transit across the OM, are likely limiting steps, see below). Localizing CglB to the cell surface is nevertheless essential because CglB is an OM lipoprotein^14, 15^; as such it would canonically be thought to insert into the periplasmic leaflet of the OM, though the prevalence of surface-exposed lipoproteins is becoming more widely acknowledged^33–35^. Indeed, Proteinase K shaving of liquid-grown cells (in which the motility complex is not active) did not affect CglB stability, suggesting that CglB is periplasmically-oriented under these conditions (**Supplementary Fig. S5A**). One possibility is that CglB surface exposure is directly regulated by the Glt OM platform. To explore this possibility, we first compared CglB levels in whole-cell samples of each respective *glt* mutant strain. While present at comparable levels in Δ*gltC, D, E, F, G, H, I,* and *J* backgrounds, cell-associated CglB was severely depleted in OM-platform mutants Δ*gltA*, Δ*gltB*, and Δ*gltK* (but not Δ*gltH*) (**Fig. 3A**). Fractionation analysis revealed that CglB was still produced by the Δ*gltA*/*B*/*K* mutants (**Fig. 3B**); however, unlike in WT cells — where CglB was detected in whole-cell and outer-membrane vesicle (OMV) fractions — CglB in these three mutant backgrounds was only recovered in culture supernatants (**Fig. 3B, Supplementary Fig. S5B**). In the Δ*gltA* and Δ*gltB* mutants, such shedding to the supernatant was not observed for GltK (**Fig. 3B**), another OM lipoprotein required for gliding motility which remains OMV-associated, nor for the intracellular protein MglA, which was detected at levels comparable to the WT strain (**Fig. 3B**); together, these data indicate that cell integrity is not affected when components of the OM platform are disrupted. Therefore, cell-association (and thus OM localization) of CglB depends on GltA, GltB, and GltK.

**Figure 3:**
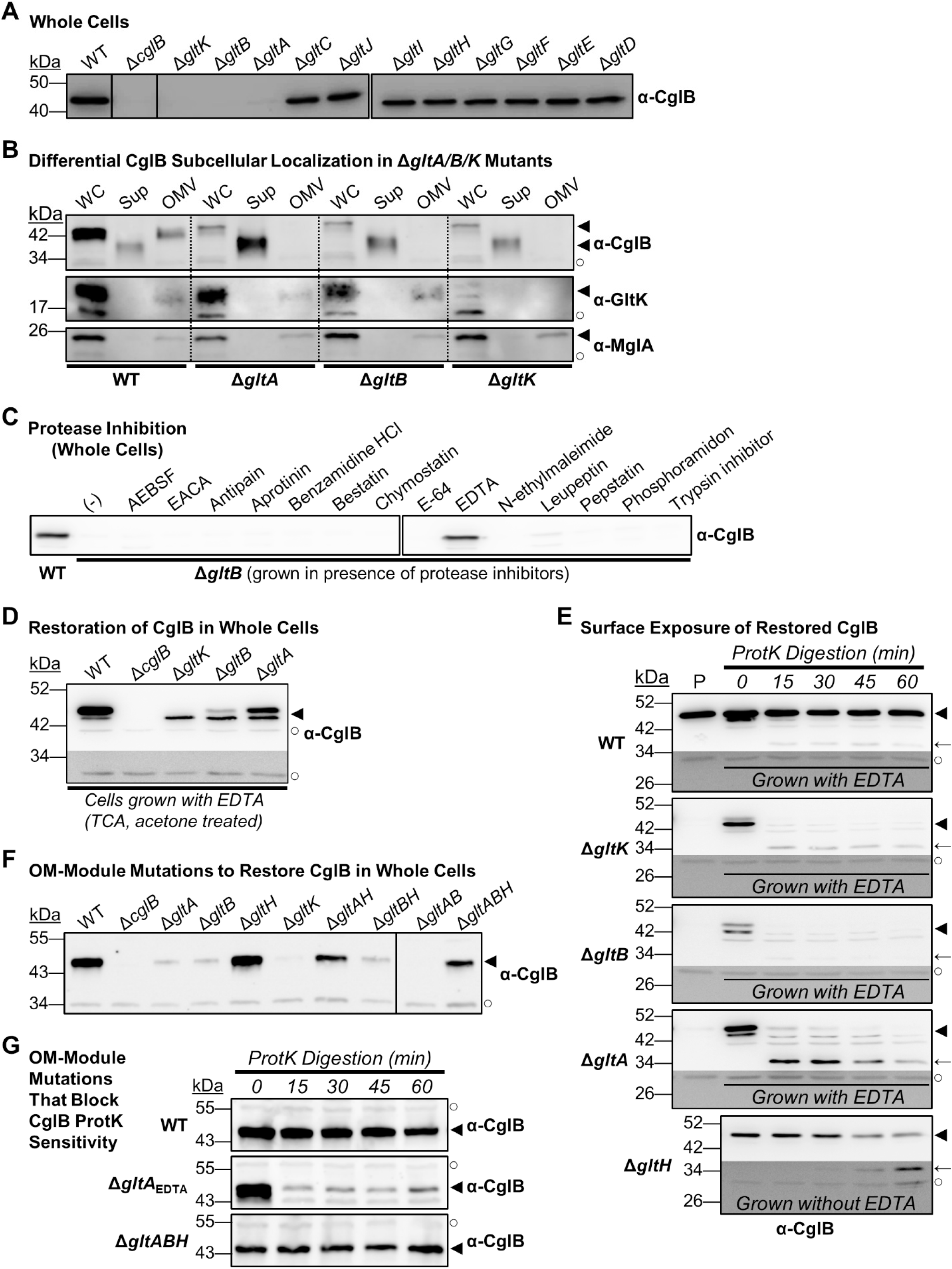
CglB Secretion is Mediated by the Glt OM Platform α-CglB Western blots from: **A)** Whole-cell extracts from different Δ*glt* mutants. Non-adjacent lanes on the blot are separated by vertical black lines. White space separates two distinct blots processed at the same time. **B)** Fractionated samples containing whole cells (WC), supernatants (Sup), and outer-membrane vesicles (OMV) from various genetic backgrounds. Detection of the gliding motility OM lipoprotein GltK was added as a control, with the protein only detected in WC and OMV samples, showing that the various mutations do not affect OM integrity, with the supernatant localization in this instance being specific to CglB. MglA is a cytoplasmic protein added as a control to show that cell lysis is negligible and does not account for the presence of CglB in supernatants. Legend: ◄, full-length protein; ○, loading control (non-specific protein band labelled by the respective pAb). **C)** Whole-cell extracts from Δ*gltB* cells grown in the presence of different protease inhibitors. White space separates two distinct blots from the same experiment. **D)** Restoration of cell-associated CglB in EDTA-grown Δ*gltK*/*B*/*A* cells. Different ratios of the slower- and faster-migrating bands were observed depending on the mutation of the specific strain. Lower, darker zones on each blot correspond to sections of the same blot image for which the contrast has been increased to highlight lower-intensity protein bands. Legend: ◄, full-length CglB; ○, loading control (non-specific protein band labelled by α-CglB pAb). **E)** Protein samples from cells resuspended in TPM buffer and digested with exogenous Proteinase K. Aliquots of the digestion mixture were removed at 15-min intervals and TCA-precipitated to stop digestion. “P” and “P_EDTA_” denote lanes containing the untreated parent strain and the parent strain grown in the presence of EDTA, respectively. Lower, darker zones on each blot correspond to sections of the same blot image for which the contrast has been increased to highlight lower-intensity protein bands. Legend: ◄, full-length CglB; ←, CglB degradation band; ○, loading control (non-specific protein band labelled by α-CglB antibody). See also **Supplementary Fig. S5A**. **F)** Whole-cell extracts from different combinations of Δ*glt* OM-module mutations in the same strain. Non-adjacent lanes on the blot are separated by vertical black lines. Legend: ◄, full-length CglB; ○, loading control (non-specific protein band labelled by α-CglB pAb). **G)** Protein samples from cells resuspended in TPM buffer and digested with exogenous Proteinase K. Aliquots of the digestion mixture were removed at 15-min intervals and TCA-precipitated to stop digestion. Legend: ◄, full-length CglB; ○, loading control (non-specific protein band labelled by α-CglB pAb).

Supernatant-localized CglB migrates faster than cell-associated CglB via SDS-PAGE (in both whole-cell and OMV samples) under equivalent denaturing conditions (**Fig. 3B**), suggesting that supernatant CglB is of reduced molecular weight and may have been proteolytically cleaved. Further support for proteolytic processing of CglB was provided via mass spectrometry analysis of tryptic peptides obtained from immunoprecipitated supernatant CglB, which revealed that the first 76 N-terminal residues were unaccounted for (**Supplementary Fig. S5C**). Our efforts at N-terminal sequencing of supernatant CglB were inconclusive, and as such we were unable to determine the specific identity of the amino acids at the new N-terminus of the truncated protein. Thus CglB may be cleaved by a protease prior to its release into the supernatant. To test this hypothesis, we screened the effect of various protease inhibitors for their capacity at restoring CglB localization to the cell envelope. Growth in the presence of EDTA restored cell-associated CglB in this background (**Fig. 3C**). Similarly, EDTA also restored cell-associated CglB in the Δ*gltK* and Δ*gltA* mutants (**Fig. 3D**; see below for discussion of doublet band). Resuspension of EDTA-grown Δ*gltB* cells in EDTA-free buffer resulted in the resumption of CglB release from the cells, indicating that CglB restoration is not permanent, consistent with a protease inhibition effect (**Supplementary Fig. S5D**). Since EDTA chelates divalent cations, CglB release from Δ*gltK*/*B*/*A* cells would be consistent with the activity of a metalloprotease. While the involvement of a putative protease in CglB surface shedding awaits further clarification, the results suggest that interactions between CglB, GltA, GltB, and GltK regulate CglB cell-surface association.

To determine the subcellular site of CglB proteolytic processing in the Δ*gltA*, *B* and *K* mutants, we took advantage of the fact that EDTA blocks proteolysis and probed surface-exposure of CglB via the Proteinase K shaving assay. In marked contrast to WT cells and in the presence of EDTA, cell-associated CglB (**Fig. 3D**) was immediately digested by Proteinase K in the Δ*gltK*, Δ*gltB*, and Δ*gltA* backgrounds (**Fig. 3E**, see below for discussion of doublet band). Interestingly, upon digestion of full-length CglB in Δ*gltA* cells, there was an immediate appearance of a ∼34 kDa CglB degradation product that persisted throughout the time course, suggesting that it was protected from further digestion. This protection required GltK and GltB as the ∼34 kDa product was almost undetectable in these backgrounds (discussed further below). GltH is not as central as GltA and GltB with respect to surface retention of CglB because CglB remains cell-associated in the *gltH* mutant (**Fig. 3A**). Nevertheless, CglB also shows increased sensitivity to Proteinase K in the absence of GltH (with the appearance and steady accumulation of an ∼34 kDa Proteinase K-resistant band), suggesting that a fraction of the CglB population is also surface exposed (**Fig. 3E**). These results thus indicate that CglB becomes surface-exposed in the absence of individual OM-platform components GltA, B, K, and to a lesser extent GltH (see Discussion).

### The Glt OM proteins are required for CglB exposure at the cell surface

Two hypotheses could explain the cell-surface protease sensitivity of CglB in Δ*gltA*/*B*/*K*/*H* cells: (i) CglB accesses the cell surface via an as-yet unknown system and interacts with the Glt OM platform proteins, which shield it from the action of the putative surface metalloprotease. Alternatively, (ii) the Glt OM platform proteins are directly responsible for CglB cell-surface exposure through a regulated process that becomes constitutive as soon as one of its components (i.e. GltA, B, K, and to a lesser extent H) is removed.

To discriminate between these hypotheses, we probed cell-association of CglB in Δ*gltABH* triple-mutant cells lacking any of the β-barrel components of the OM platform. The level of cell-associated CglB in these triple-mutant cells was markedly restored (although not fully to WT levels, possibly due to stabilization of CglB by interactions with the OM gliding proteins) (**Fig. 3F**). This rules out the first hypothesis. Using various combinations of double β-barrel component knockouts, CglB restoration was found to minimally require the simultaneous deletion of *gltA* and *gltH* (**Fig. 3F**). To examine whether the restoration of cell-associated CglB was due to a lack of surface exposure of the adhesin, we performed Proteinase K shaving of Δ*gltABH* cells. In this background, CglB was fully protected from digestion by Proteinase K, illustrating that CglB is no longer surface-exposed (**Fig. 3G**). Taken together, the above results indicate that interactions between GltA, B, K and H regulate CglB exposure at the cell surface. Importantly, full Proteinase K-protection of CglB in the Δ*gltABH* triple mutant (i) is a strong indication that these proteins play a central role in modulating CglB surface exposure, and (ii) rules out non-specific OM defects in the absence of OM platform components.

### GltK is required for CglB function in vivo

While analysing CglB restoration in EDTA-grown cells, CglB was observed to migrate as two bands in WT cells: a major slower-migrating band and a minor faster-migrating band (**Fig. 3D**). In similar conditions, only the faster-moving band was observed in the Δ*gltK* background. Thus, GltK could be involved in the formation of the slower-migrating species, a process in which it may be assisted by GltB and to a lesser extent by GltA, because the slower-migrating form accumulated to various ratios in these mutants, with almost full efficiency in the Δ*gltA* mutant (**Fig. 3D**). It is not currently clear if the two CglB forms result from a stabilization or maturation effect of GltK on CglB, but the slower-migrating CglB band constitutes the most abundant species in WT extracts (**Fig. 3D**) and thus probably represents the functional form of CglB, which we subsequently confirmed (see below).

GltK could potentially interact with CglB via Cys residues as it also possesses a high Cys ratio (9 out of 184 aa = 4.9%) relative to other Glt proteins (**Supplementary Fig. S2B**) and displays redox-dependent folding (**Fig. 4A**); but in the absence of an *in vitro* assay with purified proteins, this could not be directly tested. Nevertheless, the potential for GltK to promote CglB function *in vivo* was examined via two further experiments. First, TIRFM-analyzed clusters in Δ*gltK* cells revealed cluster behaviour analogous to that observed in Δ*cglB* cells (**Fig. 2C-G**); namely, clusters travelled directionally within Δ*gltK* cells with lifetimes equivalent to those in WT cells, but with higher trafficking frequency (**Fig. 4B, C, D**). The trafficking speed of clusters was found to be statistically faster in Δ*gltK* cells compared to WT (**Supplementary Fig. S4E**), but within the same tight range already described for Δ*cglB* cells (**Fig. 2F**). In contrast, no trafficking clusters were observed in Δ*gltA* cells (**Fig. 4B**), consistent with previous results^8^. This demonstrates that although GltA is required for cluster formation and trafficking, GltK is not (similar to CglB). Combined with the Proteinase K-susceptibility data (**Fig. 1B, C**), these results suggest that in the absence of GltK, the OM platform is assembled and conducive to AglZ-YFP trafficking but that CglB-mediated adhesion is not functional.

**Figure 4:**
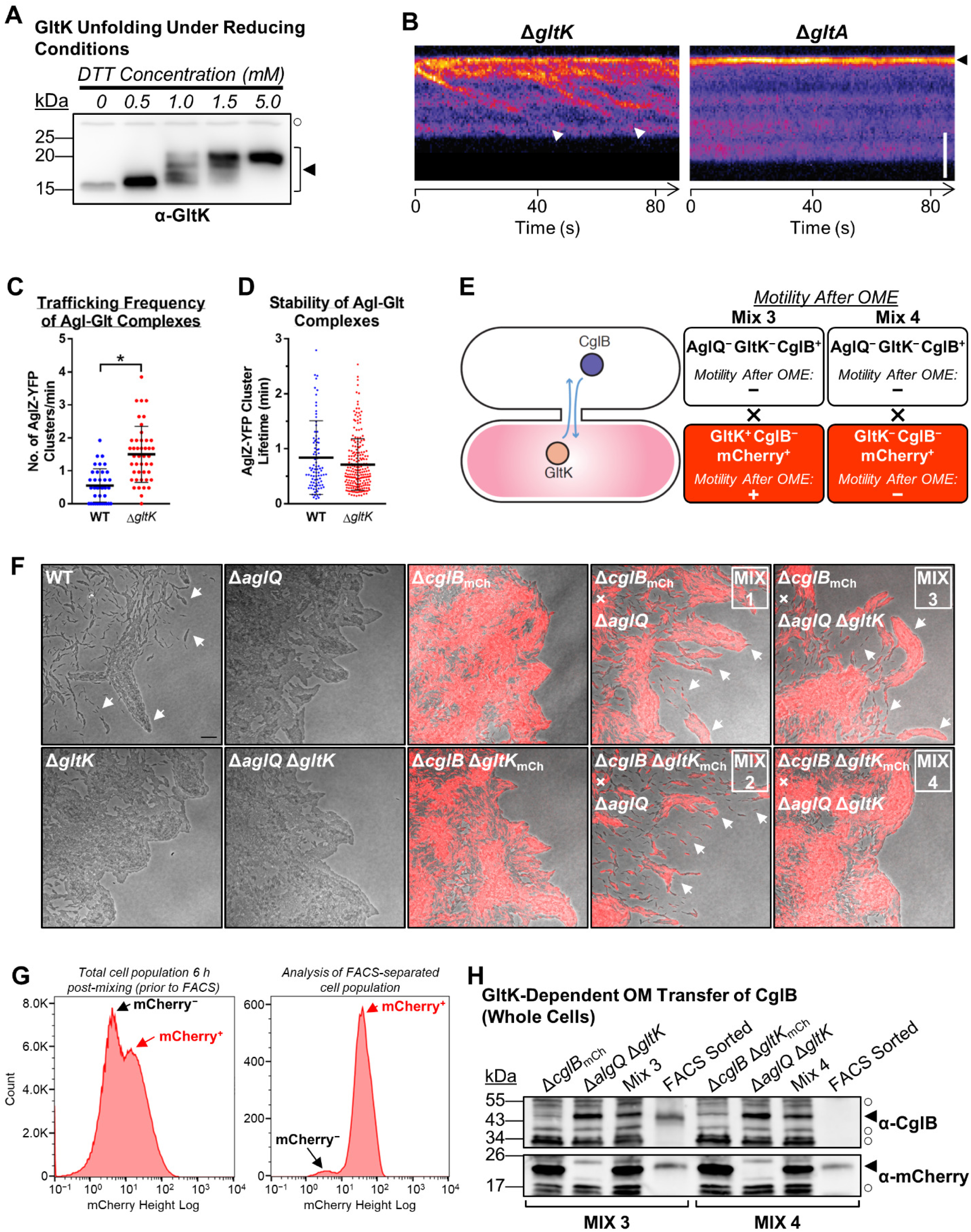
Role of GltK in CglB Function. **A)** α-GltK Western blot of WT whole-cell extracts treated with increasing concentrations of DTT to break disulphide bonds. Legend: ◄, full-length GltK; ○, loading control (non-specific protein band labelled by α-GltK pAb). See also **Supplementary Fig. S2B**. **B)** Kymograph of Δ*gltA* vs. Δ*gltK* cells showing AglZ-YFP cluster position over time. Scale bar: 2 µm. Legend: white arrowheads, AglZ-YFP clusters followed for their entire lifetime; black arrowheads, AglZ-YFP clusters followed for an incomplete lifetime. See also **Supplementary Fig. S4C**. **C)** Frequency of trafficking Agl–Glt complexes via TIRFM (of AglZ-YFP) on chitosan-coated glass surfaces in PDMS microfluidic chambers for WT (n = 37 cells) and Δ*gltK* (n = 44 cells) strains. The distribution of the two datasets are significantly different (*), as calculated via unpaired two-tailed Mann-Whitney U-test (*p* < 0.05). See also **Supplementary Fig. S4C**. **D)** Stability of trafficking Agl–Glt complexes via TIRFM (of AglZ-YFP) on chitosan-coated glass surfaces in PDMS microfluidic chambers for WT (n = 90 clusters) and Δ*gltK* (n = 223 clusters) strains. The distribution of the two datasets are not significantly different, as calculated via unpaired two-tailed Mann-Whitney U-test (*p* > 0.05). See also **Supplementary Fig. S4C**. **E)** Schematic of inter-strain mixes to study OME and the interplay between GltK and CglB. **F)** Gliding motility restoration in mCherry^+^ cells via *trans*-complementation following OME. Note that motile single cells, as well as gliding-dependent flares emanating from swarm fronts (which are dependent on collective effects of single-cell motility), are only observable when CglB is efficiently transferred (white arrows). Scale bar: 20 µm. **G)** *M. xanthus* cells from strain-mixing experiments to probe GltK-dependent CglB transfer via OME. *Left panel:* Fluorescence profile of the total cell population after 6 h of cell mixing on an agar substratum, as determined via flow cytometry. *Right panel:* FACS analysis of the mCherry^+^ cell population after sorting. **H)** Western immunoblot to detect the presence of CglB and mCherry following OME via various strain–strain mixes. Legend: ◄, full-length protein; ○, loading control (non-specific protein band labelled by respective pAb).

We next sought to genetically demonstrate that GltK permits CglB to function *in vivo*. We thus exploited the known ability of *M. xanthus* to transfer OM lipoproteins between compatible cells by a process known as OM exchange (OME)^36^. During OME, direct interactions of TraA OM receptors lead to transient OM fusion synapses between adjacent cells and efficient exchange of OM lipids and lipoproteins between contacting cells^37, 38^. In this manner, OME of CglB can efficiently rescue gliding motility when donor CglB^+^ cells are mixed with recipient CglB^−^ cells^15^. GltK (previously CglC) is also a transferable OM lipoprotein^13, 17^. Therefore, we reasoned that if Δ*gltK* cells were mixed with Δ*cglB* cells, motility rescue of the Δ*cglB* cells would only occur if bi-directional OME allows GltK transfer (from the Δ*cglB* donor cells) to activate CglB in the Δ*gltK* cells (**Fig. 4E**). To ensure that gliding motility rescue could only arise from the *trans*-complementation of the Δ*cglB* cells, an *aglQ* mutation was added to the CglB^+^ donor, thus deleting an essential component of the Agl gliding motor^11^. This motor protein cannot be transferred via OME as it is an integral IM protein. Introduction of a fluorescent mCherry marker into the CglB^−^ recipient strain allowed for (i) direct observation of motility rescue (**Fig. 4F**) and (ii) separation of mCherry^+^ cells via fluorescence-activated cell sorting (FACS) (**Fig. 4G**) for testing of CglB transfer by Western immunoblot. The Δ*aglQ* donor strain functioned as an effective donor for both CglB and GltK, ensuring that the experiment could be performed (**Fig. 4F**, Mix 1 & Mix 2). Upon mixing Δ*aglQ* Δ*gltK* double-mutant cells (GltK^−^ CglB^+^) with mCherry^+^ Δ*cglB* (GltK^+^ CglB^−^) cells, gliding motility of the mCherry^+^ cells was restored (**Fig. 4F**, Mix 3). Western immunoblotting of FACS-separated mCherry^+^ cells (**Fig. 4G**) confirmed that this gliding restoration was due to CglB transfer (**Fig. 4H**).

The prediction that GltK transfer from GltK^+^ CglB^−^ cells to GltK^−^ CglB^+^ cells is essential for CglB transfer and stimulation of GltK^+^ CglB^−^ cell motility was tested by mixing Δ*aglQ* Δ*gltK* cells (GltK^−^ CglB^+^) with an mCherry^+^ Δ*cglB* Δ*gltK* (CglB^−^ GltK^−^) strain. In this case, no motility stimulation was observed (**Fig. 4E,F**, Mix 4) nor was any CglB transferred via OME (**Fig. 4H**). The lack of CglB transfer is likely due to a defect in CglB maturation/stabilization that occurs in the absence of GltK because mixing a Δ*gltA* mutant — where CglB is also secreted but where the slower-migrating CglB form accumulates to near-WT levels (**Fig. 3D**) — with a Δ*cglB* mutant still resulted in efficient motility stimulation (GltA itself is not transferable [**Supplementary Fig. S6**]). GltK is thus required for CglB function *in vivo* by promoting the accumulation of the slower-migrating CglB band, which either occurs as GltK protects CglB from proteolysis or promotes a maturation step.

## Discussion

Previously, we demonstrated that on hard surfaces *Myxococcus* cells are propelled by directionally-transported Agl–Glt complexes that become tethered at bFAs where they exert traction forces against the underlying substratum^8^. However, while we demonstrated that the external OM Glt proteins are effectively anchored to the surface, the specific protein(s) conferring the adhesive properties, as well as the mechanism by which such adhesion is regulated to promote forward movements, remained unknown. The characterization of CglB, a protein first studied >40 years ago^17^, and its interactions with the OM Glt proteins provides a solution to these questions.

Together, the data described in our current investigation indicate that CglB exposure at the cell surface and its proper functioning is a complex process requiring the concerted actions of a host of additional proteins such as GltK, GltB, GltA, and GltH. Combining these data with previously-published findings, we propose the following model for CglB-mediated surface coupling of the Agl–Glt complex at bFAs:

i. Following its assembly at the leading cell pole, the cytoplasmic–IM–periplasmic Agl–Glt complex contacts the periplasmic face of OM gliding machinery modules, constituted by GltKBAH “loaded” with CglB, through the peptidoglycan meshwork; this initiates a right-handed helical movement of the machinery toward the lagging cell pole. No bFA is formed until the complex reaches the “ventral side” of the cell that lies in contact with the underlying substratum (**Fig. 5, *Step 1***).
ii. Once the cytoplasmic–IM–periplasmic module contacts a substratum-associated OM module, the contractile force/action of the cytoplasmic–IM–periplasmic module on the OM complex “fires” CglB into activated contact with the substratum (**Fig. 5, *Step 2***). Substratum-anchored CglB is thus connected across the OM by GltK and the associated GltABH OM proteins, interacting with other periplasmic Glt proteins^8^. This physically links the substratum and the IM motor, allowing for force transduction (**Fig. 5, *Step 3***, see below).
iii. After the power stroke of the IM constituents of the Agl–Glt machinery, the substratum-adhered OM module is displaced, resulting in displacement of the OM and peptidoglycan sacculus (**Fig. 5, *Step 4***). Displacement requires substratum-bound CglB to be turned over to allow new OM modules (containing CglB not yet bound to the substratum) to interact with the IM motor (**Fig. 5, *Step 4***) and to propel the screw-like movement of the cell (**Fig. 5, *Step 5***). Such turnover could occur by site-specific proteolysis of CglB, regulated by the GltABHK complex (**Fig. 5, *Step 3***). This hypothesis is attractive because proteolytic turnover also occurs for integrin adhesions in metazoans^39^ and de-regulation of CglB secretion triggers cell-surface cleavage of CglB at its N-terminus. However, this proteolytic event could also be a natural consequence of unregulated surface exposure of the adhesin in liquid-grown cells, with other potential adhesin-cycling mechanisms functioning instead on solid surfaces.

**Figure 5:**
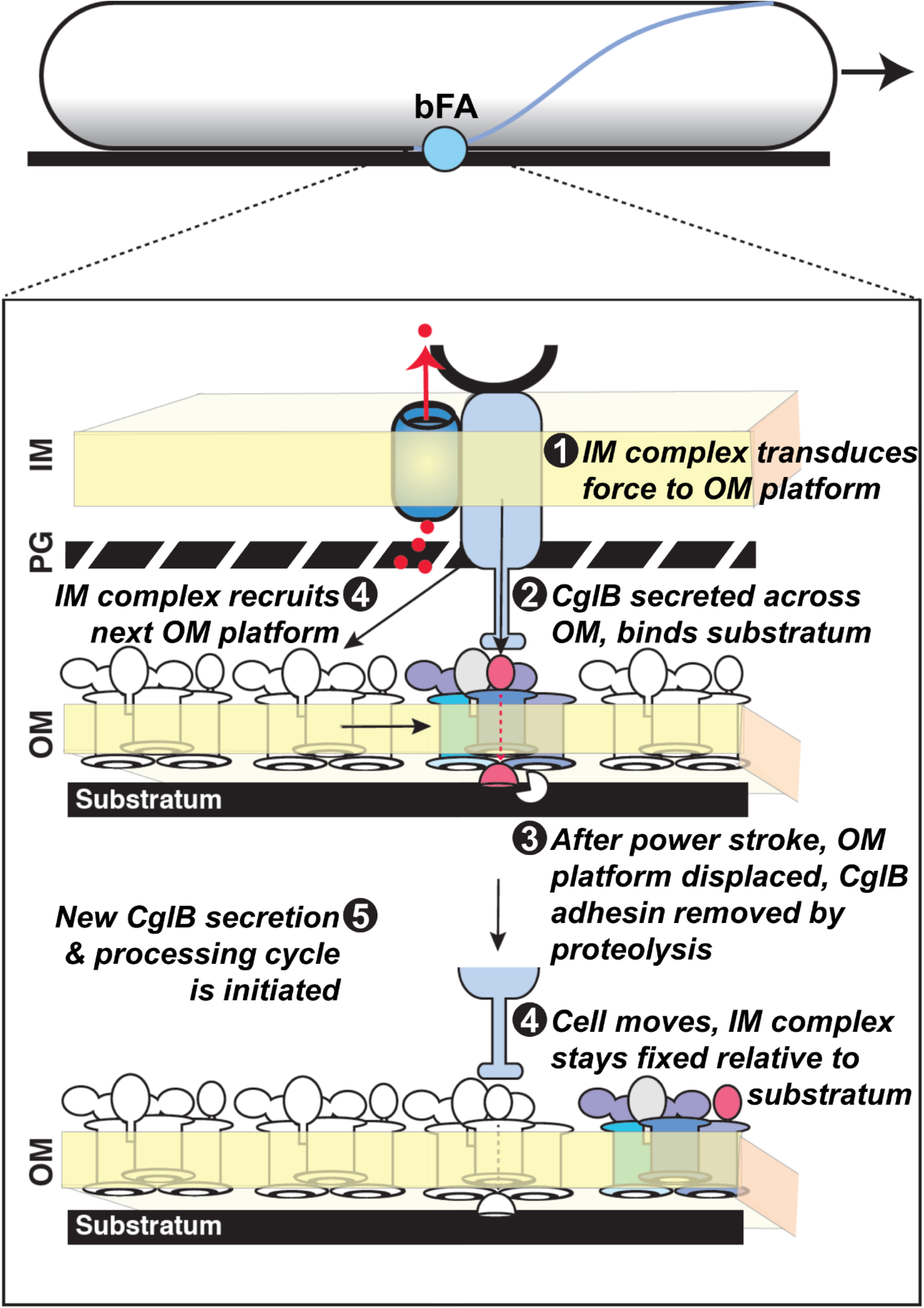
Proposed Model for Agl–Glt Complex Substratum-Tethering by CglB at bFAs. *Step 1:* At bFAs, the PMF-driven (*red dots*) mechanical action of the IM motor complex allows dynamic interactions of the IM complex with the OM platform via flexible protein periplasmic domains. *Step 2:* These domains promote localized CglB activation via GltK and secretion of CglB across the OM, requiring the OM-platform structure. CglB then interacts with the underlying substratum via its integrin-like domain, thus mediating force transduction at bFAs. *Step 3:* After the power stroke of the IM machinery, the “used” OM platform is displaced. Adhesin turnover must occur to allow the motility complex to engage new CglB-loaded OM platform complexes and mediate forward propulsion. The exact turnover mechanism remains to be identified and could involve action of a surface metalloprotease to release CglB from the OM platform. *Step 4:* Displacement of the “used” OM platform, coupled with release of the “used” adhesin, promotes forward propulsion of the cell body, while the IM complex remains spatially fixed relative to the substratum. The IM complex subsequently recruits the adjacent inactive CglB-loaded OM platform and brings it in register. *Step 5:* This again locally activates a new CglB secretion and processing cycle. This creates an OM-platform adhesin flow at bFAs, propelling the cell forward. For simplicity a single Agl–Glt complex is shown at a bFA, but it is possible that bFAs are formed by several co-localized complexes that act in concert.

Our findings assign an adhesion function to the CglB lipoprotein and provide a potential mechanism explaining the manner in which this protein accesses the cell surface. While OM lipoproteins are generally thought to be exposed on the periplasmic leaflet of the OM, surface-exposed lipoproteins have recently come to the fore in bacterial cell biology^34^. Although several scenarios have been proposed, the surface-exposure mechanism has not been solved for most of these lipoproteins^33^. Interestingly in *Escherichia coli*, RcsF, an OM lipoprotein sensor of cell-envelope stress, becomes exposed at the cell surface as it interacts with the β-barrel porin OmpA^40, 41^. For CglB, our results suggest that OM translocation could be more complex and driven by a dedicated system formed by the GltA, B, H, and K proteins. On the periplasmic leaflet of the OM, GltK promotes CglB function via a maturation or stabilization process, which is also somewhat assisted by GltA and GltB. It is not currently clear whether GltA, B, and H form a complex that secretes CglB directly or whether it regulates the secretion activity of another system. While the latter cannot be formally ruled out, the data suggests that GltH and GltA could form pores that secrete CglB redundantly, with GltH functioning as the major pore: (i) GltH is responsible for constitutive CglB secretion in the Δ*gltA, B,* and *K* mutants, and (ii) the simultaneous deletion of *gltA* and *gltH* abolishes CglB secretion. The CglB adhesin is partially accessible to Proteinase K in the absence of GltH (whereas it is fully protected in WT cells) also suggesting the existence of a minor GltH-independent route to the cell surface. Solving the actual structure and exact interactions within the OM platform is an ambitious endeavour, but it will be ultimately necessary to understand the manner in which interactions between three predicted porin-like proteins regulate CglB surface exposure.

CglB secretion in liquid-grown cells is not observed in WT cells and is likely a by-product of the sensitized genetic backgrounds. Thus in WT cells CglB surface exposure appears tightly controlled. We propose that CglB is only exposed at bFAs following dynamic interactions between the IM and OM complexes^8^. The exact protein interaction cascade has yet to be defined, but the proton-motor-associated IM-spanning proteins GltG/J have been proposed to undergo motor-driven extension/retraction cycles in a retractile mechanism that would allow cyclic interaction with the OM complex through the peptidoglycan^8^. Both proteins carry predicted TolA/TonB-like flexible periplasmic domains and globular periplasmic TonB_C domains^3, 8^. In ExbBD-TonB systems, these domains interact with TonB-box motifs in OM receptor proteins. Both OM β-barrel proteins GltB and GltA contain extended, unstructured N-terminal domains, with a potential TonB-box consensus sequence in the former^8^. Thus, the motorized IM proteins could mechanically act on the OM platform loaded with CglB, resulting in its export past the OM (through the OM platform or as-yet unidentified proteins) and activation of the adhesin upon engagement of the substratum. Conceptually, this mechanism could be compared to the firing of a gunlock cannon in which an arm (GltG/J) exerts mechanical force on a lanyard (TonB box on the Glt OM module), resulting in firing of a projectile (CglB) through the barrel of the cannon (GltABH).

We therefore propose that the gliding motility complex functions akin to a secretion apparatus, in which a trafficking unit permits the surface exposure and cycling of an adhesin when assembled at bFAs. Energy for the secretion process would come from the H^+^ gradient^11^. In bacteria, TonB-dependent transporters (TBDTs) are generally viewed as macromolecular import systems for siderophores and carbohydrates, but recently, the *Myxococcus* β-barrel TBDT Oar was found to be required for secretion of a developmental protease across the OM^42^. Thus, protein secretion may be an overlooked function of TBDTs, which should now be taken into account when studying these systems. In particular, the secretory function of the Agl–Glt system may shed light on its evolutionary origin. Previously, we demonstrated that the Agl–Glt system potentially evolved from a general class of envelope systems with functions as diverse as cellular motility and spore coat assembly^43, 44^. Bacterial genome-mining analyses indicate that deltaproteobacterial Agl–Glt-like systems may have evolved following acquisition and a dramatic expansion of a widely-conserved gammaproteobacterial “core” complex^43, 44^. Importantly, this proposed core system is constituted by seven conserved genes (encoding AglQ, R, S, GltC, D, E, and G homologues) including one encoding a putative TBDT (AglRQS–GltG) and OM proteins in the periplasmic leaflet (GltD and E). The original function of the core complex is not known but its composition could be sufficient for protein secretion. This scenario is hypothetical but the results reported herein demonstrate that *Myxococcus* gliding motility and protein secretion are intertwined; this appears to be a common theme across diverse bacterial motility mechanisms as exemplified by flagellar motility (Type-III secretion), twitching motility (Type-IV pili/Type-II secretion), and Bacteroidetes gliding motility^45, 46^. Bacteroidetes gliding presents an interesting parallel: a Type-IX secretion system secretes a gliding motility adhesin (SprB) which becomes propelled along the cell surface by the gliding machinery^47–50^. In fact, the Type-IX secretion system may be directly embedded into the Bacteroidetes gliding machinery; thus although they are evolutionarily distant, Bacteroidetes and myxobacterial gliding motility may follow similar operational principles.

Finally, the discovery that CglB acts as an essential adhesin of the gliding motility complex further highlights functional parallels between bFA and eFA mechanisms. In eukaryotic cells, the migration of surface-adhered cells via eFA-based locomotion involves the coordinated actions of a trans-envelope suite of proteins to transduce integrin-mediated cell– substratum adhesion to mechanical force and movement to propel the cell forward. Adhesion to the substratum at sites of activated integrins is regulated in multiple ways, including via cellular chaperones^51^ and surface proteases^39, 52^. Similar to eFAs, bFAs also interact with a secreted ECM^53^; herein the CglB VWA domain could potentially probe the physical properties of the substratum, akin to the scenario in eukaryotic integrins^22–24^. Thus, the fundamental requirements for integrin-coupled focal adhesion-based motility may be similar in metazoan, protozoan, and bacterial cells. These parallels could be important to further elucidate the mechanisms by which motility promotes multicellular behaviors^54^: similar to eukaryotic integrins, CglB could act as a sensor, regulating cell-cell interactions during development and predation.

## Acknowledgements

The authors would like to thank (i) Joel Selkrig and Nicholas Nickerson for valuable suggestions and troubleshooting regarding protease accessibility, (ii) Robert Fieldhouse for insightful discussions on protein modelling and evolutionary couplings, (iii) Sabrina Gauthier for cloning pSWU30-pCglB_WT_, (iv) Lotte Søgaard-Andersen for the kind gift of α-GltK, α-GltB, and α-GltA polyclonal antibodies, (v) Vincent Géli for the contribution of α-mCherry polyclonal antibodies, (vi) Alain Roussel and Renaud Vincentelli (CNRS – Aix-Marseille University, Architecture et function des macromolecules biologiques) for CglB protein used to generate pAb, and (vii) Régine Lebrun, Pascal Mansuelle, and Rémy Puppo of the Mediterranean Institute of Microbiology’s Proteomics Platform for insightful discussions and sample processing for mass spectrometry. A Discovery operating grant (no. RGPIN-2016-06637) from the Natural Sciences and Engineering Research Council of Canada and a Discovery Award (2018-1400) from the Banting Research Foundation fund work in the lab of S.T.I. The former, as well as graduate studentships from the PROTEO research network, support N.Y.J.’s and F.S.’s studies. Work in the lab of T.M. is supported through a European Research Council (ERC) starting grant (DOME 261105) and a coup d’élan pour la recherche française award (2011) from the Bettencourt–Schueller foundation. Work in the J.W.S. lab was supported by a grant (CAREER PHY-0844466) from the National Science Foundation (NSF). S.T.I. was previously supported by a post-doctoral fellowship in T.M.’s group from the Canadian Institutes of Health Research and the AMIDEX excellence program of Aix-Marseille University. L.M.F. was funded by T.M.’s ERC grant and a 4^th^-year thesis fellowship from the Fondation ARC. B.P.B was supported by funds from the Glenn Centers for Aging Research and the National Institutes of Health (P50 GM071508). Work from the M.S. lab was supported by a grant from the NSF (IOS135462). None of the abovementioned funding sources had any input in the preparation of this article, or in the work described herein.

## Author Contributions

STI and TM conceived of and planned the study. STI performed bright field microscopy and stereoscopy. STI and BJB carried out epifluorescence microscopy, with fluorescent cluster analysis by TM. JBF completed TIRFM, with analysis by LMF. STI, NYJ, and FS quantified cell motility. STI and AMB performed force microscopy, with analysis by AMB and JWS. LM and GB carried out FACS experiments. GS generated phylogenetic data. STI performed protein modelling and evolutionary coupling analysis. STI, LM, and BF generated mutant constructs. DJL tested polymertropism responses, with analysis by STI and DJL. STI, LM, and NYJ performed all Proteinase K digestions and processed all Western blots. LM performed fractionation assays. BF generated anti-CglB pAb. STI and TM wrote the manuscript. STI and TM generated figures. STI, MN, MS, AGG, JWS, and TM contributed personnel and/or funding support.

## Methods

### Bacterial Cell Culture and Phenotypic Analysis

*M. xanthus* strains were cultured in CYE (10% w/v Bacto Casitone peptone, 5% w/v yeast extract, 1% w/v MgCl_2_, 10 mM MOPS [pH 7.4]) broth with shaking (220 rpm), or on CYE solidified with 1.5% agar, at 32 °C. To examine the effects of protease inhibition on CglB liberation, cells were grown in the presence of individual protease inhibitor panel constituents (Sigma, Cat.# INHIB1) at the recommended concentration: 4-(2-Aminoethyl benzenesulfonyl fluoride HCl (AEBSF) (1 mM), ε-aminocaproic acid (EACA) (5 mg/mL), antipain HCl (100 µM), aprotinin (300 nM), benzamidine HCl hydrate (2 mM), bestatin HCl (40 µM), chymostatin (50 µg/mL), E-64 (10 µM), ethylenediaminetetraacetic acid disodium dihydrate (EDTA) (1 mM), N-ethylmaleimide (500 µM), leupeptin hemisulfate (75 µM), pepstatin A (1 µM), phosphoramidon disodium salt (10 µM), and soybean trypsin inhibitor (1 µM). Cell resuspensions were done in TPM buffer (10 mM Tris-HCl, pH 7.6, 8 mM MgSO_4_, and 1 mM KH_2_PO_4_). For OME tests, strains were grown to exponential phase in 20 mL CYE, then resuspended in TPM buffer to an OD_600_ of 5.0. For strain-mixing experiments, cell resuspensions were mixed together at a 1:1 ratio. Samples were then spotted (10 µL) on a CTT 1.5% agar plate and incubated (32 °C, 48 h) prior to microscopy analysis and/or harvest. All strains and plasmids used are listed in **Supplementary Table S1**.

### Polymertropism Testing

Aspect ratio (AR) vs. time experiments and analyses were adapted from previous work^21^ and were conducted as described previously^19^. Briefly, *M. xanthus* cells were grown in CYE broth at 28 °C to approximately 5 × 10^8^ cells/mL, then sedimented (4000 × *g*, 10 min), followed by resuspension in CYE broth to a final concentration of 5 × 10^9^ cells/mL and inoculation (4 µL) of compressed and uncompressed square CYE agar plates. Agar in compressed plates was squeezed against the plate wall and held in position by the insertion of a length of 5.56 mm outer diameter Tygon tubing^21^. Plates were incubated at 30 °C and the colony perimeters marked after 24, 48, 72, 96, and 120 h. Upon experiment completion, the AR of each swarm at each time point was calculated by dividing the colony width by the colony height such that an elongated swarm produced an AR greater than one, while a round swarm produced an AR near-or-equal to one. Linear best-fit lines were plotted for each replicate dataset, followed by slope (AR/time) determination. Slope values were averaged for each strain, then normalized as a percentage of the AR/time for the WT strain.

### DNA Manipulations

The upstream region of *cglB* (from -213 bp), including a promoter region (from -190 bp to -141 bp) predicted by BDGP^55^, as well as *cglB* itself was amplified via PCR using Q5 high-fidelity DNA polymerase, followed by digestion of the product and plasmid pSWU30 with HindIII-HF (5’) and SacI-HF (3’), then ligation via T4 DNA ligase (all enzymes from NEB) to yield pCglB_WT_. Oligonucleotide primers for QuikChange site-directed mutagenesis were generated using PrimerX (http://bioinformatics.org/primerx/). Sequencing results were analyzed by Sequencher and/or ApE software.

### Generation of α-CglB polyclonal antibodies

CglB (lacking signal peptide) elaborating a C-terminal hexa-histidine tag (CglB_21-416_- His_6_) was purified under denaturing conditions. Fractions were collected in 50 mM Tris pH 8.0, 300 mM NaCl, 250 mM imidazole, 6 M urea and used to immunize rabbits (Eurogentec, 28-day speedy protocol). The α-CglB 1° pAb produced was then tested for specificity by using the wild-type and the Ω*cglB* strains. The α-GltA, α-GltB, and α-GltH 1° pAb were raised previously^4, 13^.

### Phylogeny and gene co-occurrence

This study explored 27 myxobacterial genomes, distributed within 3 suborders and 10 families^56–69^, in addition to 58 outgroup genomes (members from 32 non-Myxococcales Deltaproteobacteria, 4 α-, 6 β-, 9 γ-, 4 £-proteobacteria, 2 Firmicutes, and 1 Actinobacteria). Highly-conserved gapless concatenated alignment of 25 housekeeping protein sequences^66, 70^ was subjected to RAxML to build a maximum likelihood phylogenetic tree using JTT Substitution Matrix and 100 bootstrap values^71^. Sequential distribution of gliding motility genes, i.e. *agl*, *glt* (M1, G1 and G2 clusters)^4^ and *cglB*^15, 72, 73^ was identified within all 85 genomes under study using two iterations of homology searching via JackHMMER (HMMER 3.1b1 [May 2013])^74^ with an E-value cut-off of 1e^-5^, query coverage of 35%, and 35% sequence similarity. The relative distribution of gliding motility proteins was mapped to the multi-protein phylogeny using iTol v3^75^. The strip right to the phylogeny depicts the taxonomic classes (from top to bottom: Myxococcales, non-Myxococcales δ-proteobacteria, α-, β-, γ-proteobacteria, and fibrobacteres respectively).

### Tertiary structure homology detection & modelling

Pair-wise alignments were performed with EMBOSS Needle software^76^. Protein secondary structure was analyzed via Jpred4^77^, with domain detection performed using SMART^78^. Preliminary identification of structural homologues to CglB, GltK, GltB, GltA, and GltH was carried out using fold-recognition searches of the Protein Data Bank using HHpred^79^ and RaptorX^80^. To improve template–target alignments, outputs were manually curated in MODalign^81^ to minimize breaks in predicted target secondary structure motifs. Curated alignments to well-matched templates were then input into MODELLER^82^ to generate a 3° structure homology model.

CMView^83^ was used to identify intra-protein contacts between Cα positions within model structures with a minimum sequence separation of 10 amino acids. Evolutionarily-coupled amino acid positions within proteins were identified using CoinFold^84^, with the highest-stringency *L* (length of the protein) number of coupled residues extracted. Positions of coupled amino acids as well as intra-protein contacts were individually plotted in GraphPad Prism v.6, followed by tricolour overlay using ImageJ to identify overlapping positions and colour swapping in Photoshop (Adobe) to improve clarity.

### SDS-PAGE, in-gel fluorescence, and Western immunoblotting

For detection of proteins from whole cells via Western immunoblot, TPM-washed cells were sedimented and resuspended at OD_600_ 1.0 in 1× Laemmli sample buffer containing 5% β-mercaptoethanol for reducing SDS-PAGE (unless otherwise indicated). Samples were boiled (10 min), loaded (20 µL) on 10-well 1 mm-thick gels, then resolved on 10% acrylamide gels (80 V for 45 min for stacking, 120 V for 75 min for resolving), then electroblotted (100 V for 60 min) to nitrocellulose membranes. Blots were rinsed with Tris-buffered saline (TBS) buffer, blocked for 30 min at room temperature with 5% milk in TBS, then incubated rocking overnight in the 4°C cold room in 1:10 000 α-CglB, or 1:3000 α-GltH, or 1:1000 α-mCherry pAb mixture in TBS with 0.05% Tween-20 (TBS-T). The next day, blots were rinsed twice (5 min) with TBS-T, incubated with goat α-rabbit 2° antibody conjugated to HRP (1:5000) (Bio-Rad) in TBS-T at room temperature (1 h), then rinsed twice (5 min) again with TBS-T. For all detections using α-GltA (1:5000) and α-GltB (1:5000) pAb, identical processing steps were followed, but using PBS-based buffers (instead of TBS). All immunoblots were developed using the SuperSignal West Pico (Thermo) chemiluminescence substrate, captured on either a GE Imager with ImageQuant software or an Amersham Imager 600 machine.

For fractionated whole cell–supernatant–OMV samples in 1× Laemmli buffer, samples were boiled (10 min) and loaded (20 µL) on 15-well 4-20 % acrylamide precast gradient gels (Biorad). Supernatant-alone samples were similarly boiled and loaded on a cast 10% acrylamide gel. Gels were resolved at 120 V, followed by electroblotting to nitrocellulose membranes at 100 V. Immunodetection was performed with diluted polyclonal antisera as follows: α-CglB (1:10 000), α-MglA (1:5000), and α-GltK (1:5000). Detection via secondary antibody was done with goat α-rabbit mAb (1:5000) conjugated to HRP (Biorad). Immunoblots were developed using the SuperSignal West Femto (Thermo) chemiluminescence substrate, captured on GE Imager with ImageQuant software.

For analysis of AglZ-YFP in-gel fluorescence, TPM-washed cells were resuspended in 1× non-reducing Laemmli sample buffer to an OD_600_ of 4.0. Cell resuspensions were heated for 30 min (65 °C), loaded (20 µL) on an 8% SDS-PAGE gel, and resolved for 45 min at 80 V, then 75 min at 120 V. Cultures, cell resuspensions, and SDS-PAGE gels (before, during, and after resolution) were all shielded from ambient light to reduce photobleaching of the YFP moiety. Resolved gels were scanned on a Typhoon FLA9500 flat-bed imager (GE Healthcare). AglZ-YFP was excited with a 473 nm laser, with fluorescence captured using the BPB1 filter (PMT 800). Pre-stained protein ladder bands were detected via excitation with a 635 nm laser and capture using the LPR filter (PMT 800). Quantification of band fluorescence intensity was performed using ImageJ via the “plot lanes” function, followed by determination of the area under the curve. AglZ-YFP signal for each lane was normalized to the faster-migrating auto-fluorescent band in the same lane; these values were then expressed as a percentage of the signal in WT cells for a given biological replicate.

### Sample Fractionation

To separate supernatant and outer-membrane vesicle (OMV) fractions, WT, Δ*gltA*, Δ*gltB* and Δ*gltK* vegetative cells were grown in CYE medium to OD_600_ 0.7. Intact cells were first eliminated by sedimentation at 7830 rpm (10 min, RT). After addition of 1 mM PMSF, supernatants were sedimented at 125 000 × *g* (2 h, 4 °C). The resulting pellets (OMV fraction) and supernatants (soluble fractions) were then treated separately. The OMV pellets were washed with TPM, sedimented again at 125 000 × *g* (2 h, 4 °C), and then resuspended directly in 500 µL 1× Laemmli protein sample buffer. The soluble supernatant fractions were treated with TCA (10 % final concentration) for 30 min on ice and then sedimented at 11 000 rpm (1 h, 4 °C). The resulting pellets (precipitated proteins) were washed with 100% acetone, sedimented at 7830 rpm (10 min, 4 °C), and dried overnight at RT. Dried pellets were then resuspended in 1.5 mL TPM, sedimented at 15 000 rpm (30 min, 4 °C) and finally resuspended in 500 µL 1× Laemmli protein sample buffer.

For isolation of supernatant-alone samples, 50 mL CYE cultures (inoculated at OD_600_ 0.02) were grown overnight with shaking (220 rpm, 32 °C) to OD_600_ 0.2-0.3, sedimented (6000 × *g*, 15 min, 20 °C), followed by passage of supernatants through a 0.22 µm syringe filter to remove any intact cells and cell debris. Filtered supernatants were each spun through a Vivaspin 20 3-kDa cutoff column with polyethersulfone membrane (8000 × *g*, 60 min, 22 °C, fixed-angle rotor) to concentrate proteins contained therein to 1 mL final volume. Concentrated supernatants were then sedimented in a tabletop ultracentrifuge (MLA130 rotor, 120 000 × *g*, 1.25 h, 4 °C) to remove any OMVs or cell debris left in the samples. Clarified 1 mL supernatant samples were transferred to microfuge tubes and treated with 200 µL 100% TCA to precipitate the proteins. Tubes were heated at 65 °C for 5 min, then spun (16 300 × *g*, 20 min, RT) to sediment precipitate. TCA-precipitated pellets were washed with 1 mL acetone, sedimented (16 300 × *g*, 20 min, RT), followed by supernatant aspiration. Protein pellets were left uncapped in the chemical hood overnight to ensure evaporation of acetone. Pellets were resuspended in 500 µL 2× Laemmli sample buffer lacking reducing agent, then diluted to 1× with ddH_2_O.

### Immunoprecipitation & mass spectrometry

Cells of Δ*gltB* from 100 mL CYE cultures were sedimented (4000 × *g*, 24 °C, 15 min). Supernatants were decanted, pooled, and passed through a 0.2 µm syringe filter. Filtered supernatant was then concentrated using four Vivaspin20 columns (10 kDa cutoff) (Sartorius), spun at 8000 × *g* (20 °C) in a fixed-angle centrifuge, with the supernatant concentrated to the dead volume limit of each column. Concentrated supernatants (∼ 80 µL each) were subsequently pooled, and diluted 1:2 with filter-sterilized 1× PBS (binding buffer) to equilibrate sample pH. Separately, a single 1 mL Pierce Protein A column (Thermo) per each pooled supernatant was equilibrated in filter-sterilized binding buffer at room temperature as per the manufacturer’s instructions. Filtered α-CglB antiserum (1 mL) was sedimented in a microfuge to remove remnant cells and/or debris (4000 × *g*, 5 min), diluted 1:1 with binding buffer, then sedimented at 12 000 × *g* to clarify the sample as binding buffer addition may have resulted in lipoprotein precipitation. The Protein A column was primed by passage of 5 mL binding buffer. To bind antibody to the column, the 2 mL of diluted antiserum was added to the top of the column and allowed to drip through, followed by washing with 15 mL of binding buffer to remove unbound pAb. The ∼ 960 µL of supernatant concentrate was added to the top of the column and allowed to distribute throughout the resin bed at room temperature (60 min). The column was then again washed with 15 mL binding buffer. To elute bound pAb (and any associated proteins) from the column, 5 mL of elution buffer (0.1 M glycine, pH to 2.5 with HCl) was added.

To analyze the protein content of the pull-down, 500 µL of column eluate was concentrated in a microfuge using a Vivaspin500 column (10 kDa cutoff) to a dead volume of ∼20 µL, then diluted 1:1 with 2× reducing Laemmli sample buffer. Samples (20 µL) were run into the stacking gel via SDS-PAGE (80 V, 13 min). Gel bands stained with SimplyBlue Safestain were excised from the stacking portion of the gel and the proteins digested by trypsin or Endoproteinase Glu-C. Liquid chromatography coupled to tandem mass spectrometry (LC-MS/MS) analyses were performed on a Q-Exactive plus mass spectrometer (ThermoFisher Scientific) by staff at the Proteomics Platform of the Mediterranean Institute of Microbiology (Marseille). Processing of the spectra for protein identification was performed with PROTEOME DISCOVERER software (Thermo Scientific, versions 1.4.0.288 and 2.1.0.81).

### Proteinase K surface digestion

Cells were resuspended in TPM at OD_600_ 2.0, followed by addition of 200 µg/mL Proteinase K and a brief vortex pulse to mix. An aliquot (50 µL) was immediately removed at t = 0 and placed into a tube containing 5 µL of 100% trichloroacetic acid (TCA). Digestion mixtures were incubated at room temperature on a rocker platform, with aliquots removed every 15 min and placed into respective pre-aliquoted tubes of TCA. Upon removal of digestion reaction aliquots, TCA-containing sample tubes were heated at 65 °C for 5 min, chilled on ice, then sedimented at 14 000 × *g* (5 min). Following supernatant removal, precipitated protein pellets were washed via resuspension in 500 µL of 100% acetone. Samples were then sedimented as before (14 000 × *g*, 5 min), followed by careful aspiration of the supernatants. Tubes were left uncapped overnight in the fume hood to promote evaporation of residual acetone, followed by storage at −80 °C until needed. Precipitated protein pellets were resuspended in 50 µL 1× Laemmli sample buffer (with reducing agent as indicated) and analyzed via SDS-PAGE and Western immunoblot.

### Motility and fluorescence analysis

For phase-contrast and fluorescence microscopy on agar pads, cells from exponentially-growing cultures were sedimented and resuspended in TPM buffer to OD_600_ 5.0, spotted (5 µL) on a glass coverslip, then overlaid with a pad of 1.5% agar prepared with TPM. For motility analysis, cells were left to adhere for 5 min prior to imaging at 32 °C using a TE2000-E-PFS microscope (Nikon) with a 40× objective and a CoolSNAP HQ2 camera (Photometrics) with Metamorph software (Molecular Devices). AglZ-YFP fluorescence was imaged using a monolithic aluminum microscope (homemade) equipped with a 1.49 NA/100X objective (Nikon Instruments) and imaged on an iXon DU 897 EMCCD camera (Andor Technology). Illumination was provided by a 488 nm DPSS laser (Vortran Stradus), and sample positioning was performed using a P611 three-axis nanopositioner (Physik Instrument). Instrument control was programmed in LabView (National Instruments) providing integrated control of all components. Cell gliding speeds were calculated using the MicrobeJ module for FIJI ^85^. Gliding cell montages were generated using FIJI. Kymograph panels were generated using the FIJI Kymograph Builder function. AglZ-YFP clusters were detected manually and tracked with the MTrackJ FIJI plugin. Using an R software script, the points of the AglZ-YFP cluster trajectories (x0, x1, …, xn; y0, y1, …, yn) were used to calculate the mean square displacement (MSD) at time

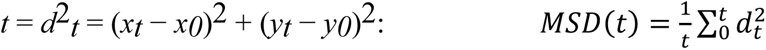

For total internal reflection fluorescence microscopy (TIRFM), imaging of real-time AglZ-YFP trafficking was performed as previously detailed in chitosan-coated polydimethylsiloxane (PDMS) microfluidic channels^8^. In brief, cells were injected into the chamber and left to adhere (30 min) without flow, with unadhered cells then removed via manual injection with TPM. TIRFM was performed on attached cells with active autofocus using an inverted microscope with 100× oil-immersion Plan-Achromat objective, atop a closed-loop piezoelectric stage. AglZ-YFP was excited with a 488-nm laser, with emission collected by the objective, through a dichroic mirror and bandpass filters, and captured by an emCCD camera. For imaging of the YFP channel in real time, 500 images were captured at 20 Hz^8^.

### Flow chamber construction and bead assay

Prior to experiments, 1 mL of *M. xanthus* DZ2 WT+AglZ-YFP and mutant DZ2 Δ*cglB*+AglZ-YFP overnight culture was grown to OD_600_ ∼0.6, sedimented (8000 rpm, 5 min), and resuspended in 400 mL TPM buffer. Flow chambers were made by combining two layers of double sided tape, a 1 mm-thick glass microscope slide, and a 100 μm-thick glass cover slip (#1.5) as previously described^86^. The tape was separated to allow a final volume of approximately 60 µL. Agarose (40 µL at 0.7%) dissolved in 6 M DMSO was injected into the chamber and allowed to sit at room temperature for 15 min. The chamber was washed with 400 µL TPM then injected with *M. xanthus* cells (60 µL) and left at room temperature to facilitate cell attachment to the agarose-coated surface for 30 min. Unattached cells were then thoroughly washed away with a total of 2 mL TPM media containing 10 mM glucose. The flow chamber was then mounted onto the microscope for imaging. For bead experiments, 1 µL of uncoated polystyrene beads (diameter 520 nm) (Bangs Laboratories) were washed and diluted in 1 mL TPM containing 10 mM glucose and injected into the flow chamber. Beads were optically trapped and placed about a third of the cell length away from the pole of the immobilized cell of interest.

### Bead tracking and video analysis

For a chosen *M. xanthus* cell (WT or mutant), 3-min movies were recorded and analyzed using a custom MATLAB tracking code. The code uses filtering mechanisms to subtract the image background from that of the cell-attached bead. Firstly, an internal MATLAB centroid function identified the *x*,*y* pixel values of the centre of the bead for each frame in the video, followed by pixel value conversion to microns. This was then used to compute motor-driven bead runs and velocities for each cell. The threshold value for a run was previously determined by disabling molecular motors and decreasing bead motion in WT cells by carefully injecting 20 μM of nigericin, a pH-gradient/proton motive force-inhibitory drug, into the mounted flow chamber. This drug concentration decreased bead velocity but not motor force production translated to the beads. In these previous experiments, 40 µM nigericin was used, leading to negligible bead motion^12^.

### Flow cytometry

Analysis of the presence of CglB in mCherry^+^ cells within a mixed population was performed with strains TM1380 (or TM1290) and TM1365, positive and negative for the mCherry marker, respectively. Fluorescence-activated cell sorting (FACS) was performed with a Bio-Rad S3E cell sorter. The blue laser (488 nm, 100 mW) was used for forward (FSC) and side scatter (SSC) signals and the green laser (561 nm, 100 mW) for excitation of mCherry. Signals were collected using the emission filter FL3 (615/25 nm). Bacterial samples in TPM buffer (maximum optical density of 0.1) were run in the low-pressure mode at ∼10,000 particles per second. The threshold on FSC was 0.15% and the voltages of the photomultipliers were 403, 299 and 722 volts for FSC, SSC and FL3, respectively. The density plots obtained (small angle scattering FSC versus wide angle scattering SSC signal) were gated on the population of interest and filtered to remove multiple events. Populations of 300,000 – 500,000 events were used and analyzed statistically using Prosort or FlowJo software. For sorting in “purity” mode, up to 60 million events were collected and at least ten tubes of 3.5 mL were used for sorting high-height peak values of mCherry fluorescence. A total volume of 35 mL was collected and pooled, followed by addition of 10% TCA to precipitate total protein content in the tube. Samples were sedimented via ultracentrifuge (164 700 × *g*, 1 h, 4 °C), then the pellet was resuspended in 500 µL of TPM, sedimented again (22 065 × *g*, 30 min, 4 °C), and later used for Western-blot analysis.

## Supplemental Information Titles and Legends

**Supplementary Figure S1:**
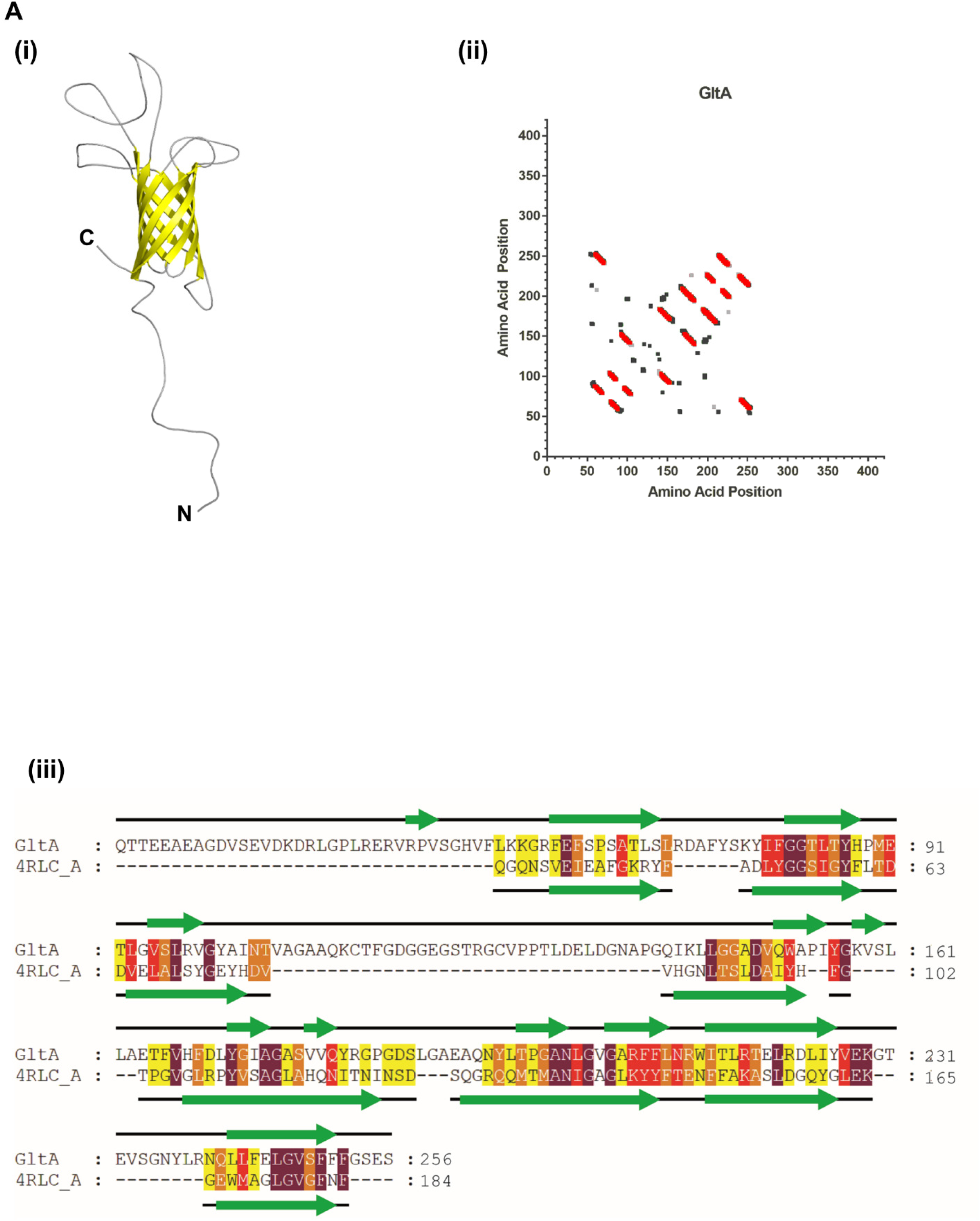
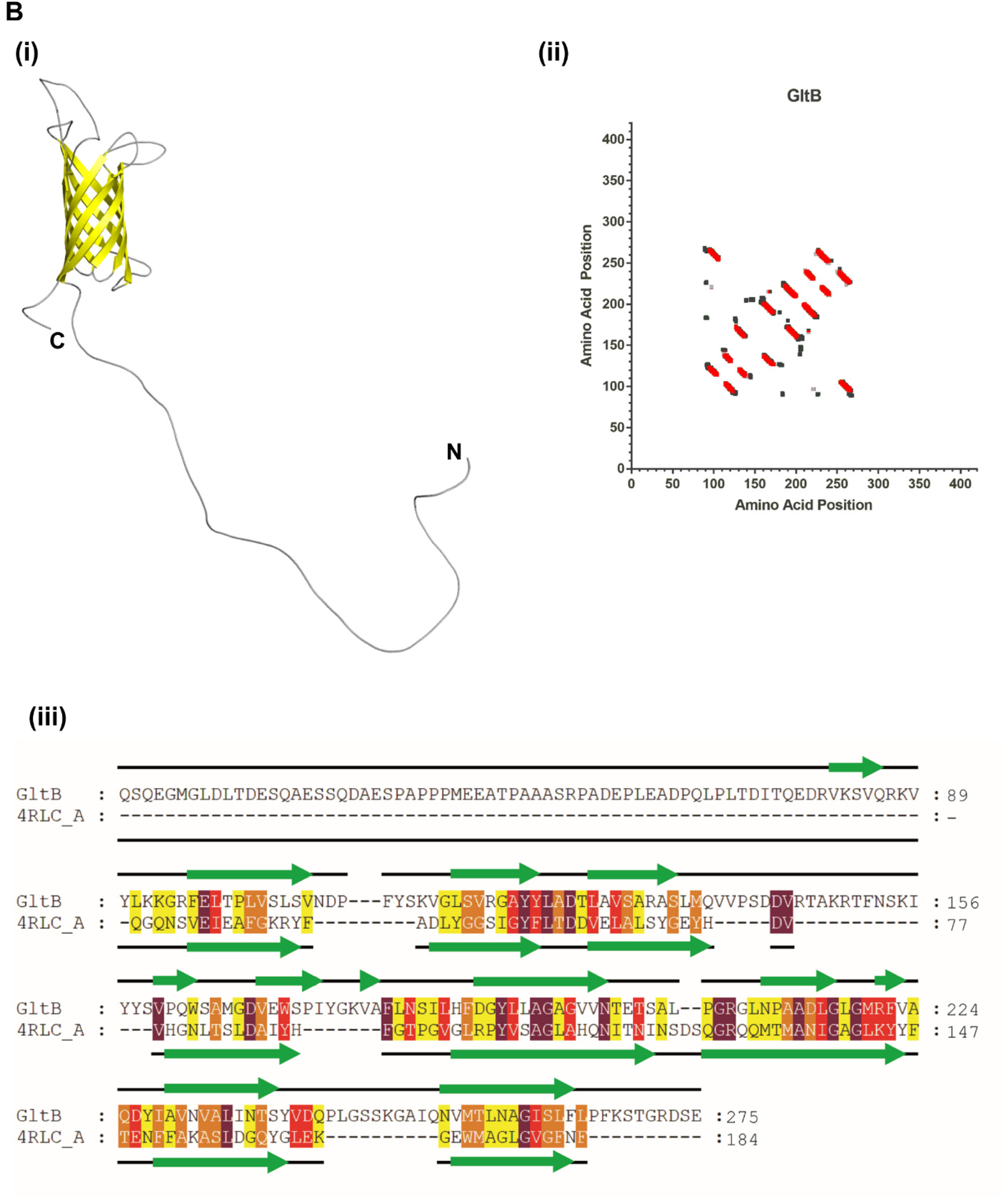
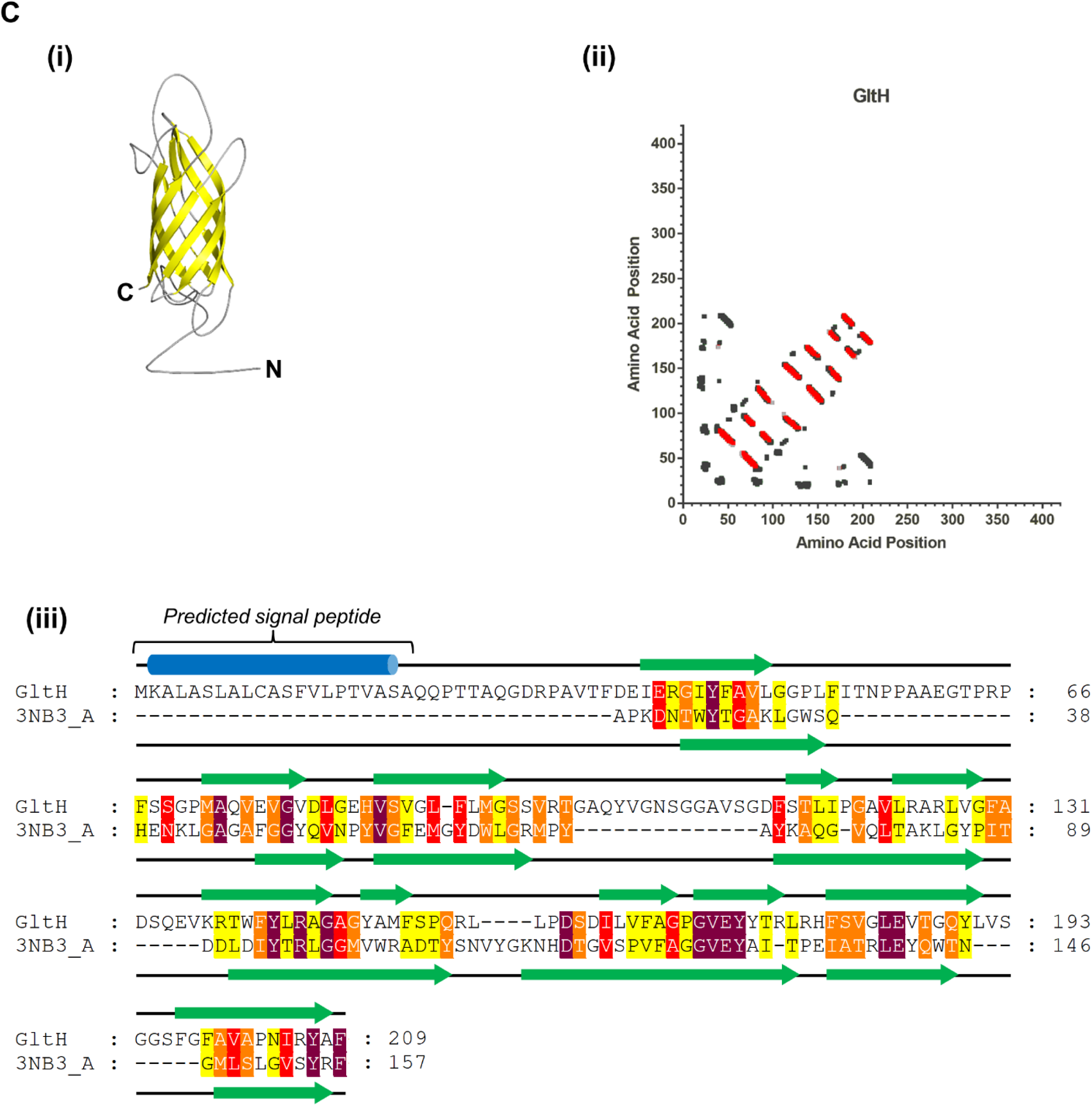
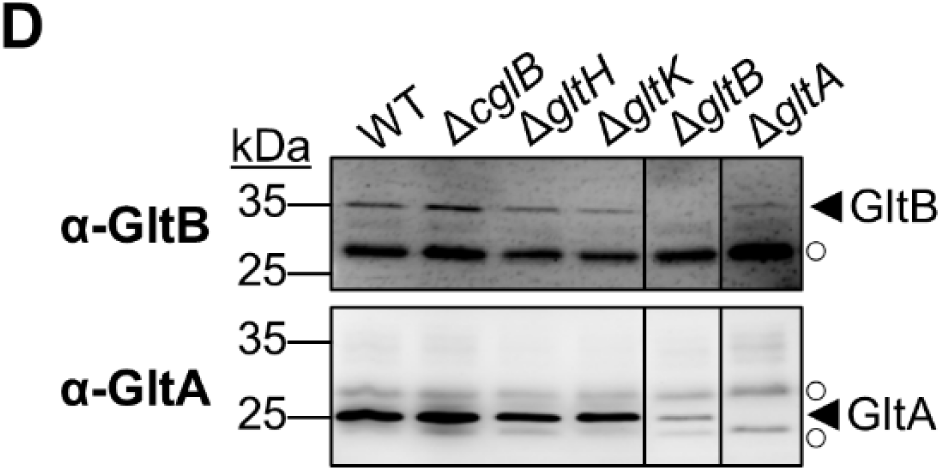
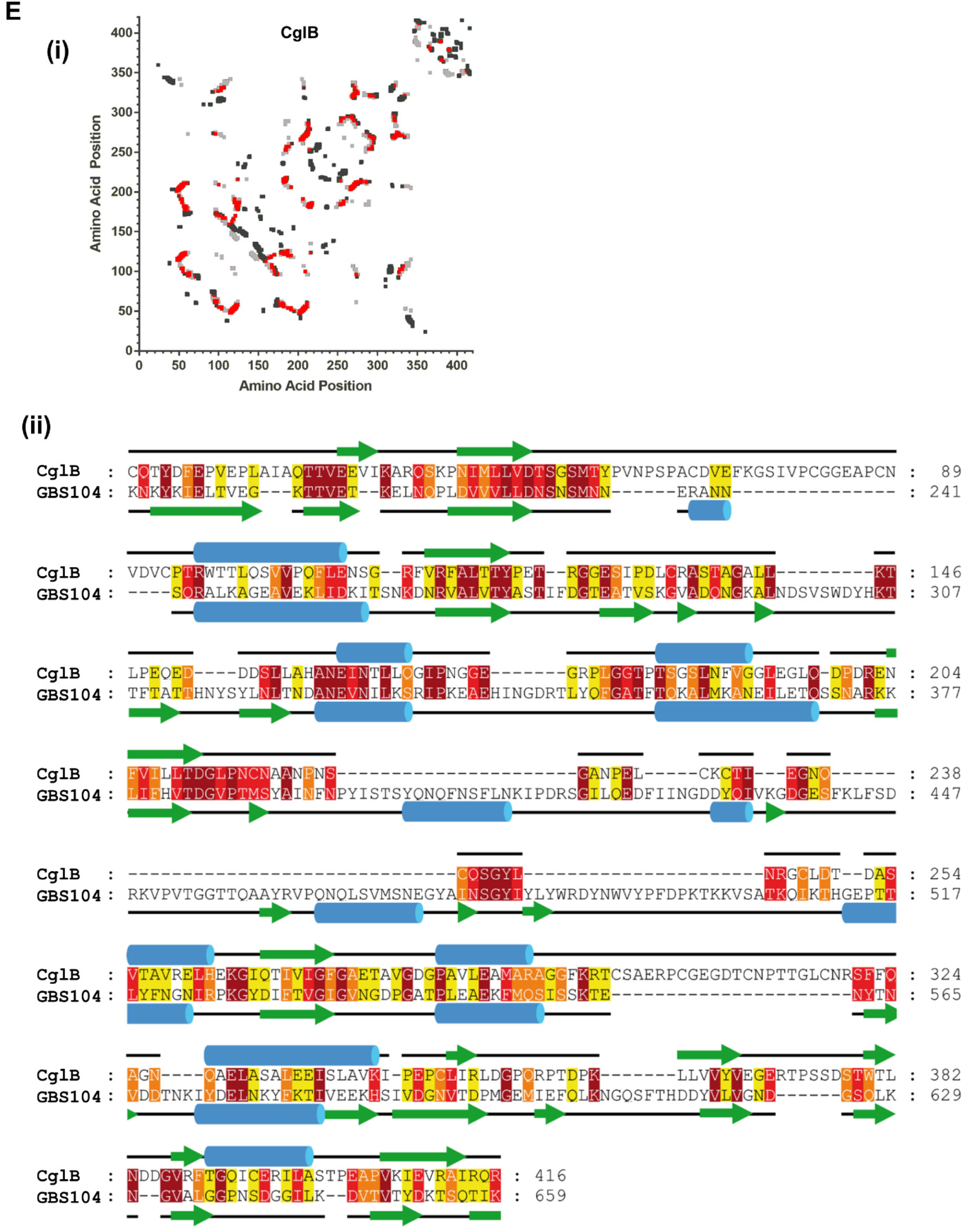
Defining the Structures of GltA, B, H, and CglB. **A)** GltA (i) tertiary structure homology model (based on the OprF β-barrel protein from *Pseudomonas aeruginosa*), (ii) overlay of intra-protein amino acid contact positions from the tertiary structure model (*black*) vs. evolutionarily-coupled amino acid positions from the primary structure (*grey*) (with conserved positions between the two datasets indicated in *red*), (iii) alignment with PDB 4RLC for MODELLER input. **B)** GltB (i) tertiary structure homology model (based on the OprF β-barrel protein from *Pseudomonas aeruginosa*), (ii) overlay of intra-protein amino acid contact positions from the tertiary structure model (*black*) vs. evolutionarily-coupled amino acid positions from the primary structure (*grey*) (with conserved positions between the two datasets indicated in *red*), (iii) alignment with PDB 4RLC for MODELLER input. **C)** GltH (i) tertiary structure homology model (based on the OmpA β-barrel protein from *Shigella flexneri*), (ii) overlay of intra-protein amino acid contact positions from the tertiary structure model (*black*) vs. evolutionarily-coupled amino acid positions from the primary structure (*grey*) (with conserved positions between the two datasets indicated in *red*), (iii) alignment with PDB 3NB3 for MODELLER input. **D)** α-GltA and α-GltB Western immunoblots from whole-cell extracts from different Δ*glt* mutants resolved via reducing SDS-PAGE. **E)** CglB (i) overlay of intra-protein amino acid contact positions from the tertiary structure model (*black*) vs. evolutionarily-coupled amino acid positions from the primary structure (*grey*) (with conserved positions between the two datasets indicated in *red*), (ii) alignment with PDB 3TXA for MODELLER input. See also Figure 2A. For the template alignments in figure panels A, B, C, E: residue colouration is based on the conservation score as determined by Jalview: maroon, 10 (out of 10); red, 9; orange, 8; yellow, 7. Conservation scores ≤ 6 have not been displayed to enhance the clarity of the figure. Secondary structure properties for each protein are indicated above/below the aligned sequence, based on the respective PDB entry and GltA/B/H/T secondary structure prediction via JPred, with β strands (*green arrows*) indicated.

**Supplementary Figure S2:**
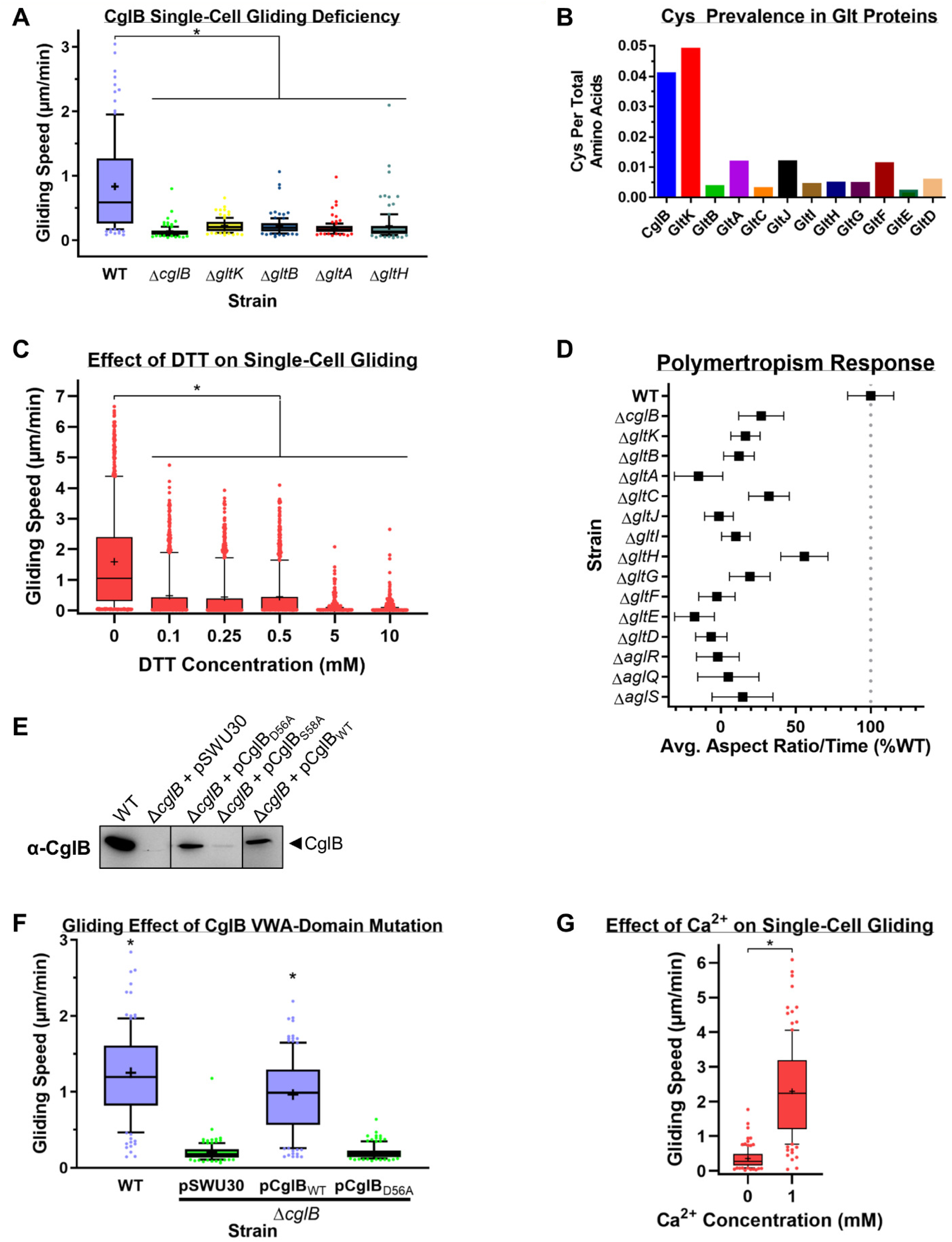
Importance of CglB to Gliding Motility and Linked Behaviours. **A)** Single-cell gliding speeds for *M. xanthus* OM-module mutant strains (n = 100 cells) on hard (1.5%) agar. The lower and upper boundaries of the boxes correspond to the 25th and 75th percentiles, respectively. The median (line through centre of boxplot) and mean (+) of each dataset are indicated. Lower and upper whiskers represent the 10^th^ and 90^th^ percentiles, respectively; data points above and below the whiskers are drawn as individual points. Asterisks denote datasets displaying statistically significant dataset differences (*p* < 0.0001) relative to WT, as determined via 1-way ANOVA with Tukey’s multiple comparisons test. **B)** Ratio of number of Cys residues per gliding motility complex protein, divided by the total number of amino acids in that protein. **C)** Single-cell gliding speeds for WT *M. xanthus* DZ2 (n = 700 cells) on hard (1.5%) agar supplemented with increasing concentrations of DTT to reduce disulphide bonds of proteins in contact with the substratum. The lower and upper boundaries of the boxes correspond to the 25th and 75th percentiles, respectively. The median (line through centre of boxplot) and mean (+) of each dataset are indicated. Lower and upper whiskers represent the 10^th^ and 90^th^ percentiles, respectively; data points above and below the whiskers are drawn as individual points. Asterisks denote datasets displaying statistically significant dataset differences (*p* < 0.0001) relative to 0 mM DTT treatment, as determined via 1-way ANOVA with Tukey’s multiple comparisons test. **D)** Polymertropism response for various *glt* and *agl* deletion mutant strains. Mean results are displayed +/− SEM, with the number of biological replicates (*n*) used to analyze each strain as follows: WT (20), Δ*cglB* (16), Δ*gltK* (19), Δ*gltB* (11), Δ*gltA* (15), Δ*gltC* (15), Δ*gltJ* (16), Δ*gltI* (15), Δ*gltH* (20) Δ*gltG* (18), Δ*gltF* (16), Δ*gltE* (14), Δ*gltD* (17), Δ*aglR* (11), Δ*aglQ* (6), Δ*aglS* (5). **E)** α-CglB Western immunoblot of CglB MIDAS motif amino acid substitution mutants from whole-cell extracts. Non-adjacent lanes on the blot are separated by vertical black lines. **F)** Single-cell gliding speeds on hard (1.5%) agar for *M. xanthus* DZ2 Δ*cglB* (n = 120 cells) complemented with CglB_WT_ or CglB_D56A_ ectopically expressed from the *attB* phage-attachment site in the chromosome. The lower and upper boundaries of the boxes correspond to the 25th and 75th percentiles, respectively. The median (line through centre of boxplot) and mean (+) of each dataset are indicated. Lower and upper whiskers represent the 10^th^ and 90^th^ percentiles, respectively; data points above and below the whiskers are drawn as individual points. Asterisks denote datasets displaying statistically significant dataset differences (*p* < 0.0001) compared to strains harbouring either the pSWU30 empty-vector control or the pCglB_D56A_ VWA-domain mutant CglB complementation construct, as determined via 1-way ANOVA with Tukey’s multiple comparisons test. **G)** Single-cell gliding speeds for *M. xanthus* DZ2 Ω*pilA* (n = 110 cells) on chitosan-coated glass in a PDMS microfluidic chamber in the presence and absence of Ca^2+^ ions. The lower and upper boundaries of the boxes correspond to the 25th and 75th percentiles, respectively. The median (line through centre of boxplot) and mean (+) of each dataset are indicated. Lower and upper whiskers represent the 10^th^ and 90^th^ percentiles, respectively; data points above and below the whiskers are drawn as individual points. Asterisk denotes datasets displaying statistically significant differences (*p* < 0.0001, as determined via two-tailed Mann-Whitney test.

**Supplementary Figure S3:**
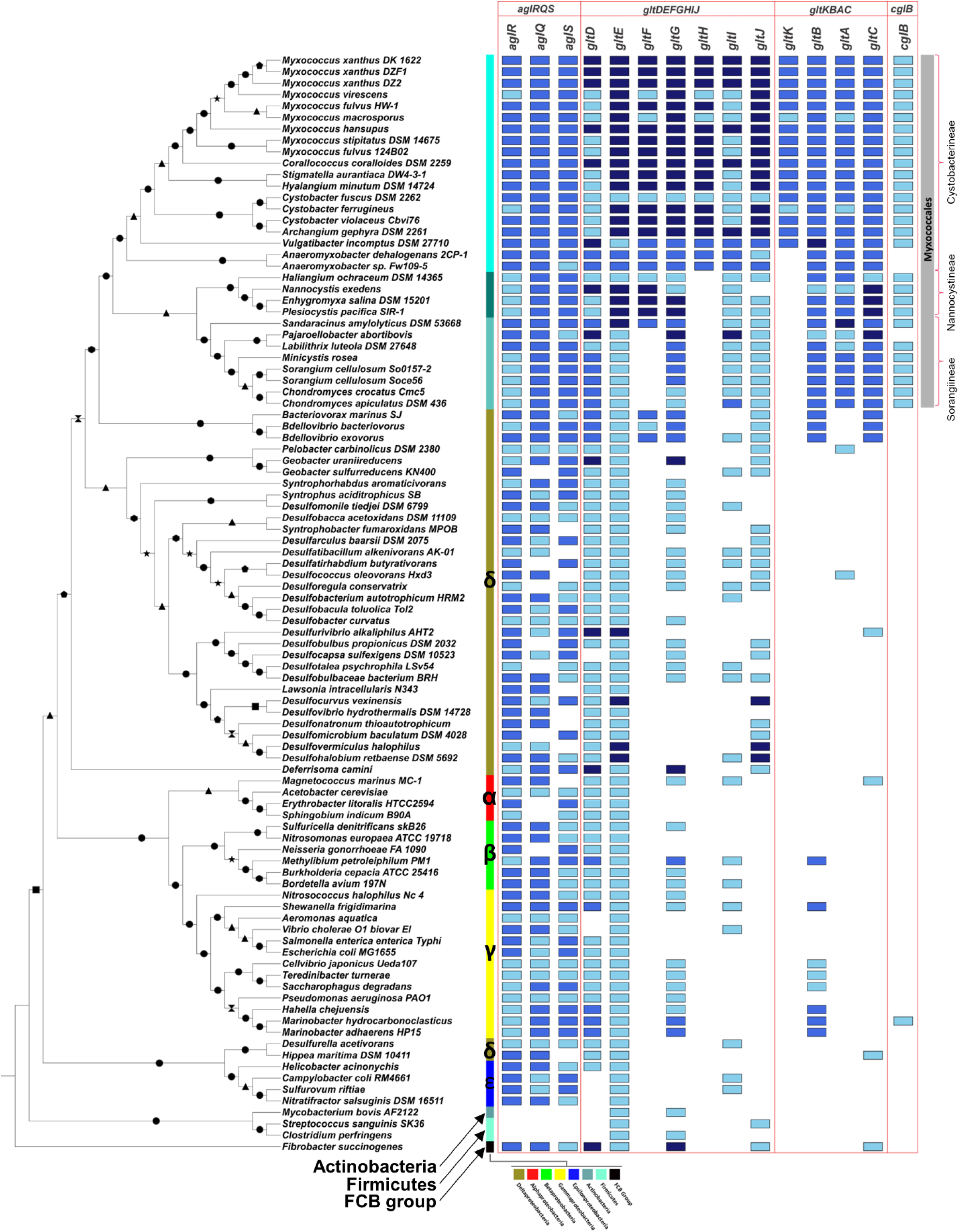
CglB Co-Occurrence and Gene Synteny in Bacteria. Taxonomic distribution and co-occurrence of *agl* and *glt* genes in bacteria. Bootstrap values at each node are indicated as follows: 100 (out of 100), e; 90-99, .._; 80-89, • ; 70-79, *; 60-69, e; 50-59, *; < 49, . Colour of gene hit indicates synteny with the G1 *gltDEFGHIJ* (*dark blue*) or G2 *gltKBAC* (*blue*) gene clusters or lack thereof (*light blue*), respectively; herein, synteny denotes a minimum of three genes in the vicinity of each other.

**Supplementary Figure S4:**
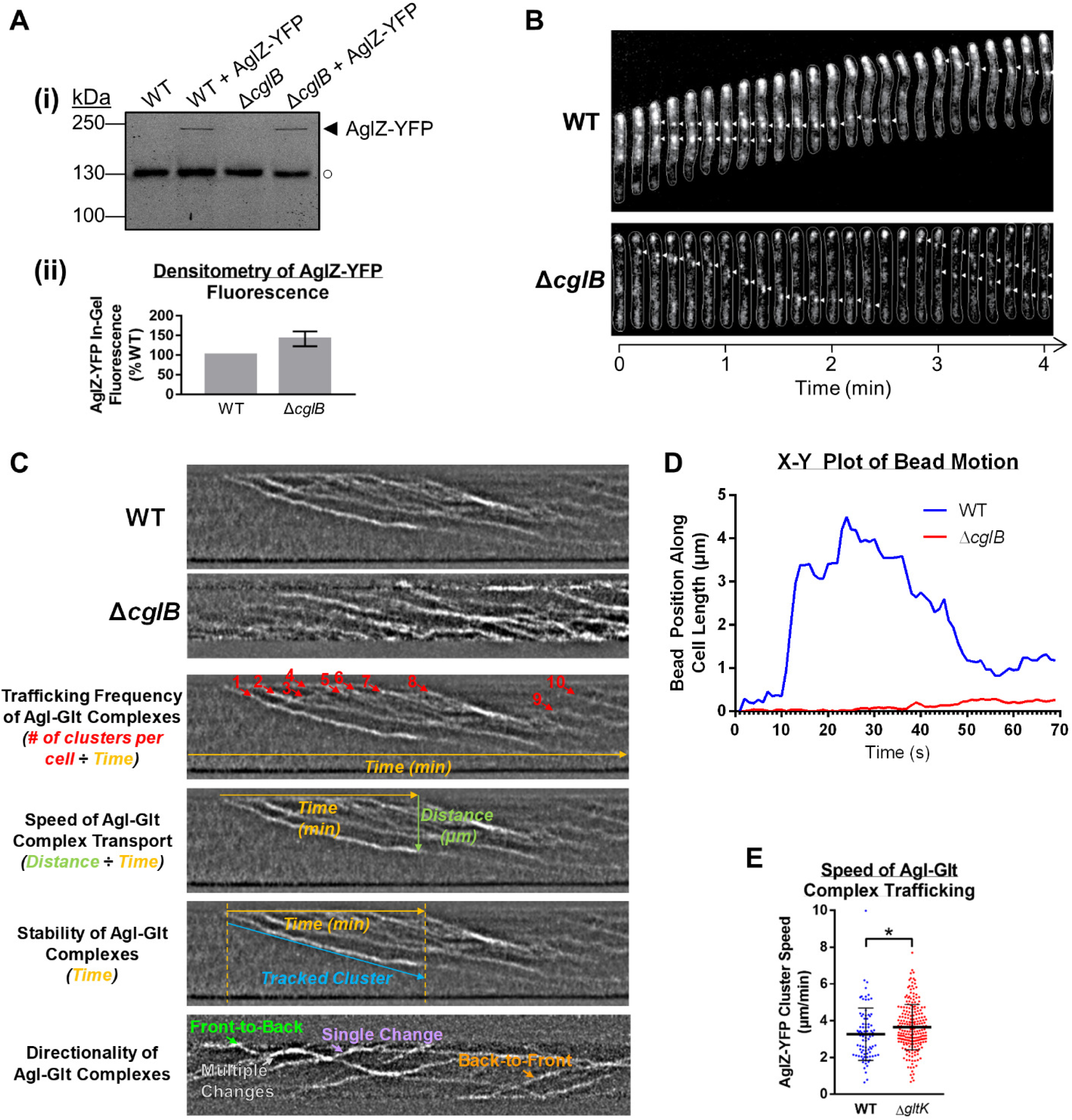
Role of CglB in Agl–Glt Complex Localization and Directed Surface Transport. **A)** In-gel fluorescence (i) scan and (ii) densitometry analysis of AglZ-YFP levels in WT vs. Δ*cglB* crude-cell lysates resolved via SDS-PAGE. Fluorescence levels were analyzed across six biological replicates and are displayed +/− SEM. Despite a higher mean value for AglZ-YFP levels in Δ*cglB* cells than in WT cells, this difference was not statistically significant, as determined via Wilcoxon signed-rank test performed relative to “100” (*p* = 0.0938). **B)** Montage of *M. xanthus* gliding time course displaying AglZ-YFP localization (white arrowheads) on hard agar via epifluorescence microscopy. Images were acquired at 10 s intervals. See also Fig. 2C. **C)** Kymograph of AglZ-YFP localization in *M. xanthus* cells on chitosan-coated PDMS microfluidic chambers via TIRFM. Arrows in *orange* denote sequential kymograph slices over time. Arrows in *cyan* indicate positions of trafficked Agl–Glt clusters in the cell. The manner in which various fluorescent cluster tracking data were obtained have been indicated in the example images. See also Fig. 2E-H. **D)** Representative time course of polystyrene bead position tracking along the length of a cell. **E)** Speed of Agl–Glt complex trafficking via TIRFM (of AglZ-YFP) on chitosan-coated glass surfaces in PDMS microfluidic chambers for WT (n = 260 clusters) and Δ*gltK* (n = 371 clusters) strains. The distribution of the two datasets are is significantly different, as calculated via unpaired two-tailed Mann-Whitney U-test (*p* < 0.05). See also Fig. 2F and **Supplementary Fig. S4C**.

**Supplementary Figure S5:**
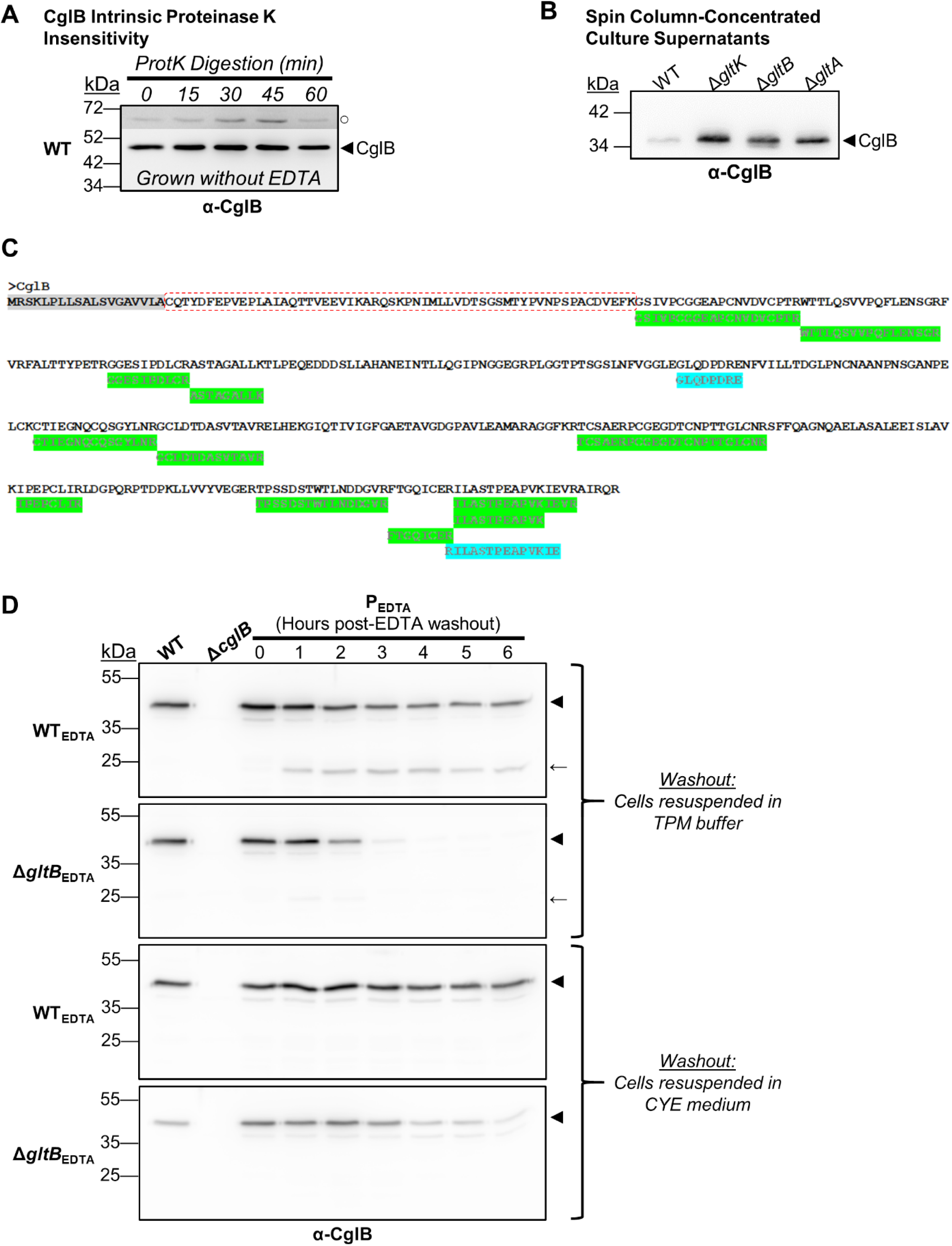
Possibility of Post-Translational Processing of CglB. **A)** Protein samples from WT cells resuspended in TPM buffer and digested with exogenous Proteinase K. Aliquots of the digestion mixture were removed at 15-min intervals and TCA-precipitated to stop digestion. The higher, darker zone on the blot corresponds to a section of the same blot image for which the contrast has been increased to highlight lower-intensity protein bands. Legend: ◄, full-length CglB; ○, loading control (non-specific protein band labelled by α-CglB pAb). **B)** Concentrated supernatant samples from Δ*glt* mutants that exhibit depleted levels of cell-associated CglB. Equivalent supernatant volumes from different strains were concentrated in a centrifuge using 3 kDa-cutoff protein concentrator columns, followed by TCA treatment of precipitate proteins. Legend: ◄, full-length CglB. See also Fig. 3B. **C)** Peptides identified via trypsin/V8 digestion and mass spectrometric analysis of immunoprecipitated CglB from Δ*gltB* culture supernatant. Legend: *grey*, predicted signal peptide; *green*, trypsin-derived peptide; *cyan*, V8-derived peptide; *red box*, N-terminal tract of CglB unaccounted for by mass spectrometry. **D)** α-CglB Western immunoblot demonstrating the resumption of CglB release from EDTA-grown cells upon transfer to an EDTA-free minimal buffer (TPM) or rich medium (CYE). Legend: ◄, full-length CglB; ←, CglB degradation band; P_EDTA_, parent strain (WT or Δ*gltB*) grown in the presence of EDTA. See also Fig. 3E.

**Supplementary Figure S6:**
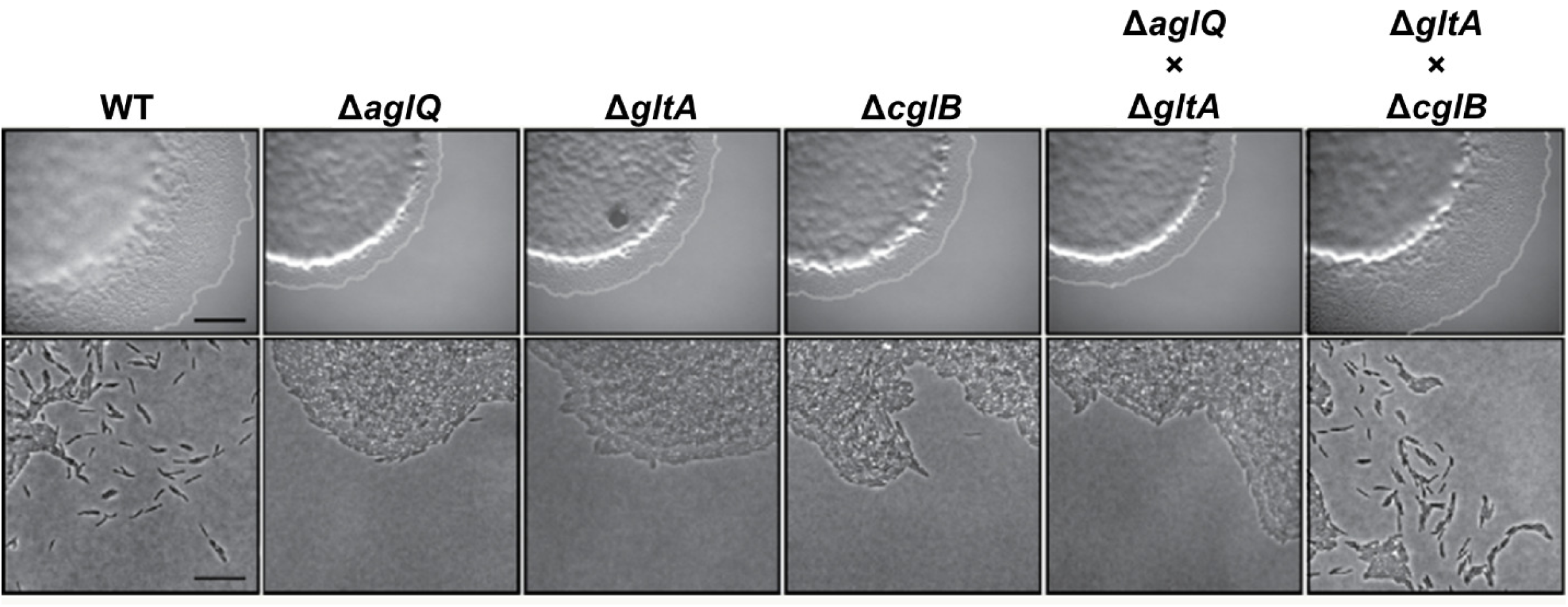
Exploration of GltK Function. Motility restoration in cells following *trans*-complementation via OME. Bottom panels (scale bar: 20 µm) are magnified views of the corresponding upper panels (scale bar: 1 mm).

**Supplementary Table S1:**
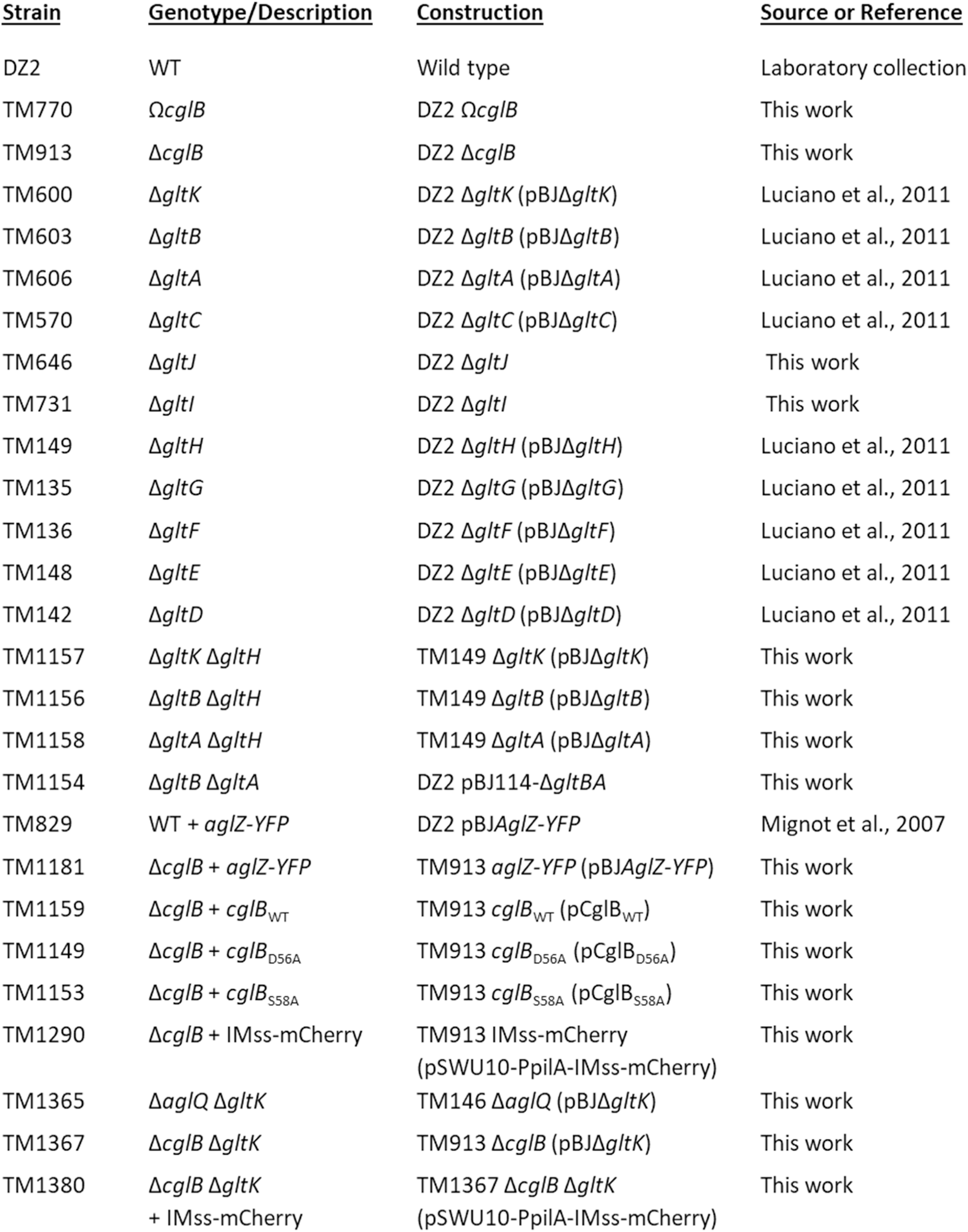
Strains of *M. xanthus* Used in this Study *Myxococcus xanthus* strains used in this study.

**Supplementary Table S2:** Fold-Recognition Hits of CglB to Various VWA-Containing Proteins Select fold-recognition hits of structural homologues in the PDB to CglB, as identified via HHpred. The strength of the hits is anchored by the strong detection of a von Willebrand Factor type A domain.

| <u>PDB Hit</u> | <u>Name</u> | <u>Probability</u> | <u>E-value</u> | <u>Score</u> | <u>Aligned</u><br><u>Coils</u> | <u>Identities</u> | <u>Similarity</u> |
| --- | --- | --- | --- | --- | --- | --- | --- |
| 1MF7_A | Integrin αM<br>(*Homo sapiens*) | 99.4 | 6.4E-14 | 123.84 | 187 | 19% | 0.191 |
| 4FX5_A | von Willebrand<br>factor type A<br>(*Catenulispora*<br>*acidiphila*) | 99.36 | 3e-13 | 140.21 | 199 | 19% | 0.161 |
| 4OKR_A | Micronemal<br>protein MIC2<br>(*Toxoplasma*<br>*gondii*) | 99.34 | 3.8E-13 | 127.97 | 180 | 20% | 0.242 |
| 4F1J_B | TRAP<br>(*Plasmodium*<br>*falciparum*) | 99.18 | 1.1E-11 | 110.16 | 178 | 17% | 0.257 |
| 3TXA_A | GBS104 tip pilin<br>(*Streptococcus*<br>*agalactiae*<br>serogroup V) | 99.13 | 7.2E-12 | 136.34 | 188 | 17% | 0.239 |

